# Mammalian UPF3A and UPF3B activate NMD independently of their EJC binding

**DOI:** 10.1101/2021.07.02.450872

**Authors:** Zhongxia Yi, René M Arvola, Sean Myers, Corinne N Dilsavor, Rabab Abu Alhasan, Bayley N Carter, Robert D Patton, Ralf Bundschuh, Guramrit Singh

**Author notes:** Correspondence, The Ohio State University, Columbus, OH 43210, USA.

## Abstract

Nonsense-mediated mRNA decay (NMD) is governed by the three conserved factors - UPF1, UPF2 and UPF3. While all three are required for NMD in yeast, UPF3B is dispensable for NMD in mammals, with its paralog UPF3A suggested to only weakly activate or even repress NMD due to its weaker binding to the exon junction complex (EJC). Here we characterize the UPF3B-dependent and -independent NMD in human cell lines knocked-out of one or both *UPF3* paralogs. We show that in human colorectal cancer HCT116 cells, EJC-mediated NMD can operate in UPF3B-dependent and -independent manner. While UPF3A is almost completely dispensable for NMD in wild-type cells, it strongly activates EJC-mediated NMD in cells lacking UPF3B. Surprisingly, this major NMD branch can operate in UPF3-independent manner questioning the idea that UPF3 is needed to bridge UPF proteins to the EJC during NMD. Complementation studies in UPF3 knockout cells further show that EJC-binding domain of UPF3 paralogs is not essential for NMD. Instead, the conserved mid domain of UPF3B, previously shown to engage with ribosome release factors, is required for its full NMD activity. Altogether, UPF3 plays a more active role in NMD than simply being a bridge between the EJC and the UPF complex.

## INTRODUCTION

Nonsense mutations present a challenging obstacle for organisms as they result in premature termination of protein translation to produce truncated proteins that can be toxic for the cell. All eukaryotes deploy a conserved mechanism called nonsense-mediated mRNA decay (NMD) to rapidly degrade mRNAs containing premature termination codons (PTCs) to limit the production of potential toxic polypeptides. NMD has gained additional importance in more complex organisms as normal mutation-free mRNAs take advantage of the NMD machinery to regulate their expression (reviewed in (He & Jacobson, 2015; Karousis & Mühlemann, 2019; Kishor et al., 2019; Kurosaki et al., 2019)). For example, in mammalian cells, ∼10% of transcriptomes can be regulated by NMD (Mendell et al., 2004; Wittmann et al., 2006). The key task for the NMD machinery is to differentiate premature translation termination from normal translation termination on both pathogenic as well as natural mRNAs that are degraded by this pathway. How the NMD machinery makes such a discrimination remains to be completely understood.

NMD depends on a set of core factors - UPF1, UPF2 and UPF3 that are conserved throughout eukaryotes. When translation terminates prematurely and much upstream of normal 3′-untranslated region (3′UTR) and polyA-tail, UPF factors can recognize such termination events as premature via mechanisms that have been conceptualized into two possible (non-mutually exclusive) models. One model suggests that termination in an altered 3′UTR context can compromise normal termination promoting interaction between release factors eRF3/eRF1 and the polyA-tail binding protein (PABP) (Amrani et al., 2004; Behm-Ansmant et al., 2007; Eberle et al., 2008; Ivanov et al., 2008; Peixeiro et al., 2012; Singh et al., 2008). Instead, UPF1 can engage with eRFs and initiate NMD (Kashima et al., 2006). According to the other model, longer 3′UTRs of NMD-targeted mRNAs may serve as a distinction between normal and premature termination. By recruiting more UPF1, the central NMD activator that can non-specifically bind RNA in a length-dependent manner, longer 3′UTRs increase the likelihood of UPF1 engagement with terminating ribosome (Hogg & Goff, 2010). While the majority of available evidence points to a more direct role for UPF1 in engaging with release factors and terminating ribosome (Ivanov et al., 2008; Kashima et al., 2006; Singh et al., 2008), a recent study shows that UPF3B (a UPF3 paralog) has a direct involvement in termination reaction in human cell extracts (Neu-Yilik et al., 2017). Nevertheless, the precise order of events and the mechanistic details of UPF functions at individual steps during premature termination remain poorly understood.

In mammalian cells, the NMD pathway has become more complex as it is tightly linked to pre-mRNA splicing via the exon junction complex (EJC), which has gained significant importance for NMD activation. The EJC is deposited on the mRNA exon-exon junctions during splicing and is exported along with the mRNAs to the cytoplasm where they are stripped-off mRNAs by translating ribosomes (reviewed in (Boehm & Gehring, 2016; Hir et al., 2016; Woodward et al., 2017)). However, when PTCs lead to early translation termination, one or more EJCs that remain bound downstream of a terminated ribosome can stimulate NMD. As UPF3 has evolved to directly interact with the EJC, the presence of EJC-UPF3-UPF2 complex in 3′UTRs can promote UPF1 activation and premature termination via either of the two NMD models. Notably, in these models UPF3 is mainly viewed as a bridge between the UPF and EJC proteins (Chamieh et al., 2008). However, the functional relevance of such a bridging function, or if UPF3-EJC interaction serves another role, remains to be seen.

While all three UPF proteins are essential for NMD in yeast (Celik et al., 2017; He et al., 1997), UPF3 appears to have become less important for NMD in more complex organisms and some NMD can proceed even in its absence (reviewed in (Yi et al., 2021)). Unlike UPF1 or UPF2, a complete loss of UPF3 in *Drosophila* does not affect viability and has only a modest effect on NMD (Avery et al., 2011). In mammals, there exist two UPF3 paralogs, UPF3A and UPF3B, and available evidence suggest that UPF3B provides the main UPF3 activity due to its better EJC binding ability (Kunz et al., 2006). UPF3A can function as a weak NMD activator and can help compensate for UPF3B function (Chan et al., 2009). However, a recent study has suggested that UPF3A may primarily function as an NMD repressor, potentially by sequestering UPF2 away from NMD complexes via its strong UPF2 binding but weaker EJC binding (Shum et al., 2016). Surprisingly, despite being the dominant NMD activating UPF3 paralog, UPF3B knockout mice are largely normal albeit with some neurological defects (Huang et al., 2011, 2018). Similarly, UPF3B inactivating mutations in humans are non-lethal although they cause intellectual disability (Laumonnier et al., 2010; Tarpey et al., 2007) and are associated with neurodevelopmental disorders such as autism spectrum disorders and schizophrenia (Addington et al., 2011; Lynch et al., 2012; Xu et al., 2013). These observations suggest that while UPF3B is important for key biological processes, its effects on NMD are likely to be peripheral since a total loss of NMD is lethal in vertebrates (Medghalchi et al., 2001; Weischenfeldt et al., 2008). Previous studies in human cell lines and mice models have shown that UPF3B is not required for NMD of several mRNAs, indicating that there exists a UPF3B-independent NMD pathway (Chan et al., 2007; Gehring et al., 2005; Huang et al., 2011). How NMD can function in the absence of UPF3 and how prevalent is such UPF3-independent NMD remains largely unknown.

UPF3B function in NMD might be further affected by specific EJC compositions. Our previous work has demonstrated that EJC composition is heterogenous and, during different phases of mRNA lifecycle, EJC associates with a distinct set of peripheral factors (Mabin et al., 2018). EJC co-factor RNPS1 does not co-exist in the same complex with another key EJC factor CASC3. Mass spectrometry of RNPS1 and CASC3 containing EJCs showed that CASC3 but not RNPS1 preferentially associates with UPF3B (Mabin et al., 2018). Consistent with this observation, a recent report found a much-reduced EJC-UPF3B association in CASC3 knockout HEK293 cells (Gerbracht et al., 2020). Together, these observations suggest a link between EJC composition and UPF3B-mediated NMD. The contribution of such a link to NMD and its underlying molecular basis remains to be fully understood.

Here, we created UPF3B knockout human cell lines with CRISPR-Cas9 to study NMD in the presence and absence of UPF3B and to understand the relative flux through the UPF3B-dependent and UPF3B-independent branches of NMD. We find that most transcripts with 3′UTR EJCs can undergo NMD in both UPF3B dependent and independent manner. In the absence of UPF3B, and only under such conditions, UPF3A becomes responsible for a significant portion of UPF3B-independent NMD. We find that while CASC3-containing EJC can moderately affect UPF3-mediated NMD, it does not appear to be the major determinant for efficient UPF3-dependent NMD. Surprisingly, our comparative analysis of UPF3A and UPF3B functions in NMD suggest that UPF3 proteins remain potent NMD activators even without their ability to bind EJC, hinting that another UPF3 function, such as its modulation of translation termination reaction, may be its primary and a more conserved mode of activating NMD.

## RESULTS

### UPF3B is required but not necessary for EJC-mediated NMD

To study UPF3B-independent NMD, we used CRISPR-Cas9 based gene editing to generate two independent *UPF3B* knockout (3B^KO^) alleles in human colorectal carcinoma HCT116 cells, a near diploid cell line with only one copy of *UPF3B*. In the first approach, we deleted ∼8 kilobase genomic region of *UPF3B* locus that spans exons 1-4 and encodes the UPF2-binding domain of UPF3B (3B^KO#1^; Figure 1A). We reason that the loss of this key functional domain will lead to a complete loss of UPF3B function during NMD. As expected, in 3B^KO#1^ cells, a smaller protein deleted of amino acids 20-155 is expressed and that too only at ∼12% of the full-length protein in wild-type cells (Figure 1B). For the 3B^KO#2^ allele, we used homology-directed repair to insert immediately downstream of the *UPF3B* start codon a puromycin resistance marker followed by a polyadenylation signal that would terminate transcription and prevent expression of the downstream sequence (Figure 1A). The resulting puromycin resistant 3B^KO#2^ cells completely lack UPF3B (Figure 1B). A qPCR survey in the two knockout cell lines revealed that mRNA levels of several previously characterized NMD-regulated genes are similarly upregulated whereas a subset of these genes remain largely unchanged in the two cell lines (Figure S1A). Thus, the HCT116 UPF3B knockout cells represent an appropriate model to study UPF3B-dependent and -independent NMD.

**Figure 1.**
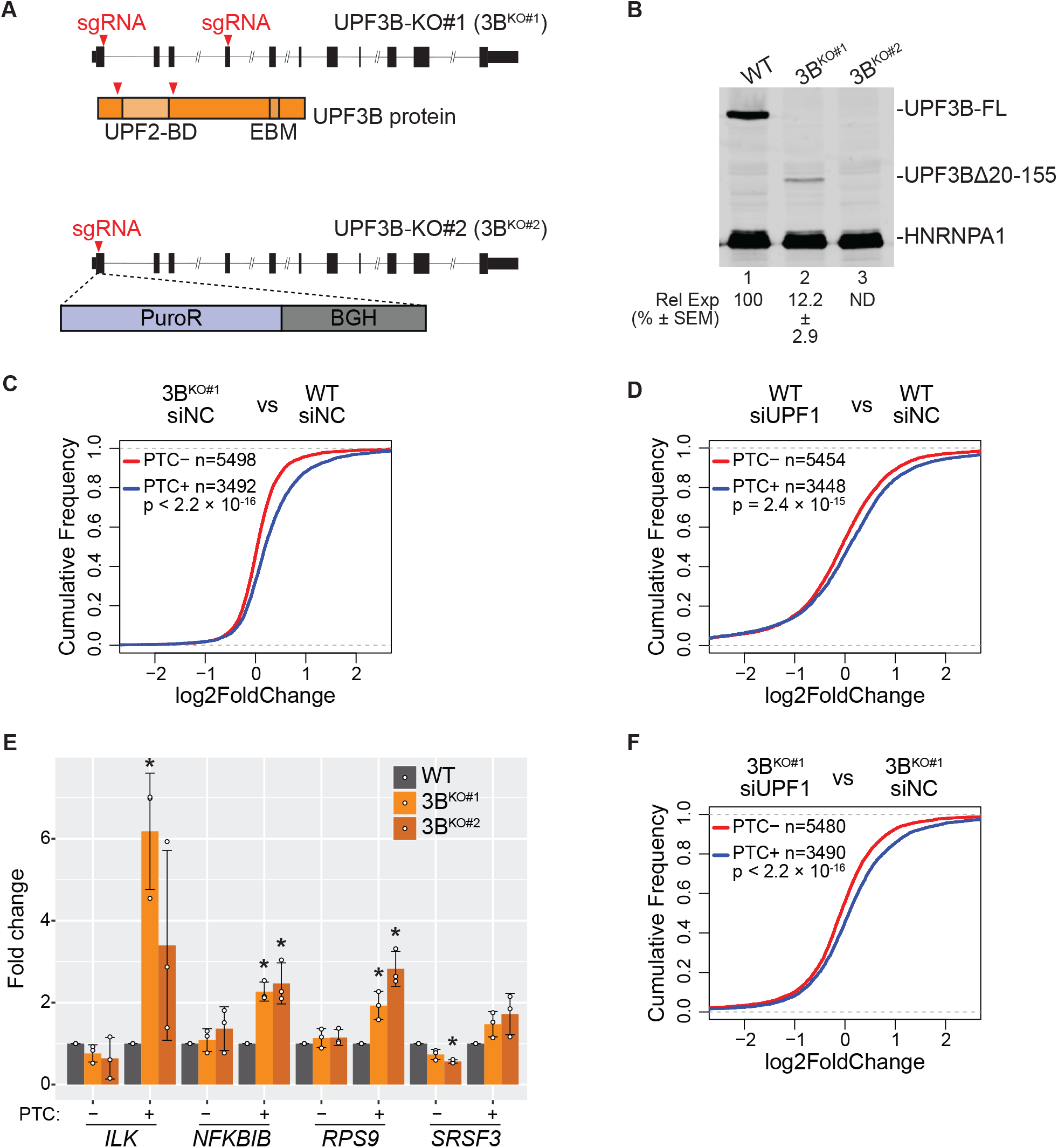
Loss of *UPF3B* in human cells affects EJC-mediated NMD. A. Schematic of *UPF3B* knockout (UPF3B-KO) strategies using CRISPR-Cas9. *UPF3B* locus is in black where rectangles represent exons and horizontal line denotes introns; coding region is shown as wider rectangles. Red arrowheads represent guide RNA targeting sites. In 3B^KO#1^ (top), two guide RNAs delete the UPF2 binding domain (UPF2-BD) of UPF3B protein coding region as shown. In 3B^KO#2^ (bottom), a donor template is used to insert puromycin resistant gene (PuroR) and bovine growth hormone (BGH) polyadenylation signal at the cut site. B. Immunoblot of wild-type (WT) and UPF3B-KO cell lines showing levels of proteins on the right. In 3B^KO#1^, a smaller UPF3B protein with deletion of amino acids 20-155 (UPF3BΔ20-155) is expressed. Relative Expression (Rel Exp) of this deletion protein as compared to the full-length WT protein along with standard error of mean (SEM) are indicated below lane 2. UPF3B antibody recognizes antigen outside the deleted region in 3B^KO#1^. HNRNPA1 is used as a loading control. C-D. Cumulative Distribution Function (CDF) plots of PTC+ isoforms and PTC-isoforms from same set of genes. X-axis represents fold change in, (C) 3B^KO#1^ versus WT cells each with control knockdown (siNC), (D) UPF1 knockdown (siUPF1) versus negative control knockdown (siNC) in WT cells. Number of transcripts in each set (n) and p-value from Kolmogorov-Smirnov (KS) test comparing the two distributions are shown. E. Bar plots from isoform specific RT-qPCR analysis showing average fold change (y-axis) of PTC+ and PTC-isoforms from genes indicated on the bottom in WT and the two 3B^KO^ cells identified in the legend on the top right. For each isoform, levels in knockout cells are compared to the levels in WT cells (set to 1). Relative levels from each replicate are shown by white circles. Error bars indicate standard error of means. The asterisk (*) represents p<0.05 in t-test with null hypothesis of true mean being 1 (n=3). F. Cumulative Distribution Function (CDF) plots of PTC+ isoforms and PTC-isoforms from same set of genes. X-axis represents fold change in UPF1 knockdown (siUPF1) versus control knockdown (siNC) in 3B^KO#1^ cells. Number of transcripts in each set (n) and p-value from Kolmogorov-Smirnov (KS) test comparing the two distributions are shown.

To identify transcriptome-wide targets of UPF3B-dependent and -independent NMD branches, we performed RNA-Seq from wild-type (WT) and 3B^KO#1^ HCT116 cells transfected with either control siRNA (siNC) or UPF1 targeting siRNA (siUPF1) (Figure S1B). The mRNAs upregulated in UPF3B knockout cells as compared to WT cells can be considered UPF3B-dependent NMD targets. Since UPF1 should be required for all NMD, mRNAs upregulated after UPF1 knockdown (KD) in UPF3B knockout cells can be considered as UPF3B-independent NMD targets. We quantified gene expression at mRNA isoform level in these RNA-seq samples and carried out differential expression analyses. As expected after loss of an mRNA repressive factor such as UPF3B, the number of upregulated transcripts in 3B^KO#1^ cells as compared to WT cells is much higher as compared to those downregulated (Figure S1C). However, the effects on the transcriptome after UPF3B loss are smaller as compared to the UPF1 knockdown (Figures S1D, E), suggesting a more restricted role of UPF3B than UPF1 in gene regulation.

Short- or long-term depletion of gene regulatory proteins such as UPF factors can cause indirect changes in gene expression (Tani et al., 2012). Indeed, we observe a similar number of up- and down-regulated transcripts after 48-hour UPF1 knockdown in HCT116 cells (Figure S1E). To minimize the impact of such indirect effects on transcriptome-wide quantification of NMD factor contributions to the pathway, we focused on a specific class of genes that produce two types of transcript isoforms, one with an exon-exon junction ≥50 nucleotides downstream of a stop codon (PTC+ transcripts) and one that lacks this well-known NMD-inducing feature (PTC-transcripts). Any change in NMD is expected to alter only the PTC+ isoforms whereas any indirect effects on gene expression are expected to similarly impact both the PTC+ and PTC-isoforms. A comparison of transcript isoform levels in 3B^KO#1^ versus control cells show a significant upregulation of PTC+ isoforms over PTC-isoforms (Figure 1C), similar to the trend observed in UPF1-KD versus control cells (Figure 1D). We confirmed the specific and significant (in most cases) upregulation of PTC+ isoforms as compared to the PTC-isoforms of several genes in UPF3B-KO cells via a qPCR assay (Figure 1E and Figure S1F). These results suggest that UPF3B is required for the efficient downregulation of EJC-dependent NMD targets in HCT116 cells.

We next examined the extent to which the EJC-mediated NMD pathway remains functional in our UPF3B knockout cells. If EJC-mediated NMD can still occur in the absence of UPF3B, we expect further upregulation of PTC+ isoforms as compared to PTC-isoforms when the NMD pathway is further compromised by UPF1 depletion in UPF3B knockout cells. In line with this prediction, we observe a significant global upregulation of PTC+ transcripts as compared to their PTC-counterparts after UPF1 knockdown in 3B^KO#1^ cells (Figure 1F), suggesting that EJC-mediated NMD can still occur in the absence of UPF3B.

In addition to EJC downstream of a stop codon, NMD can also be induced by long 3′UTRs (Eberle et al., 2008; Hogg & Goff, 2010; Singh et al., 2008). To test if UPF3B is also required for NMD of mRNAs with extended 3′UTRs, we divided transcripts based on their 3′UTR length into three groups each with a similar number of transcripts. As expected, UPF1 knockdown causes a significant upregulation of the group of transcripts with the longest 3′UTRs as compared to those with medium or short 3′UTRs (Figure S1G). Surprisingly, UPF3B knockout shows a negligible effect on the relative abundance of long 3′UTR-containing transcripts (Figure S1H). Therefore, while UPF3B is important but not essential for the EJC-induced NMD in HCT116 cells, it likely plays an insignificant role in long 3′UTR-mediated NMD in these cells.

### UPF3A replaces UPF3B in EJC-UPF complexes in UPF3B knockout cells

We next sought to address how can EJC-dependent NMD operate in human cells in the absence of UPF3B, which is widely believed to act as a bridge between the UPF proteins and the downstream EJC. To investigate the NMD complexes that assemble in the presence or absence of UPF3B, we used CRISPR-Cas9 gene editing to insert a FLAG affinity tag-encoding sequence immediately downstream of the start codon at the endogenous *UPF1* locus in both WT and UPF3B knockout cells. This facilitates FLAG immunoprecipitation (IP) of the UPF1-containing complexes from the two cell lines. We find that while in wild-type cells, UPF1 mainly associates with UPF3B and only minimally with its paralog UPF3A, in UPF3B knockout cells, UPF1-UPF3A association is dramatically enhanced (Figure 2A). Importantly, the enhanced UPF1-UPF3A association is independent of RNA. As expected based on the previous observations (Chan et al., 2009; Tarpey et al., 2007), UPF3A is upregulated 3.5-fold in UPF3B knockout cells (Figure 2B). Notably, in RNA-Seq data from 3B^KO#1^ cells, *UPF3A* mRNA shows 1.8-fold increase. Thus, overall increase in UPF3A in UPF3B knockout cells likely occurs both at the mRNA and protein level. Similar to its enhanced association with the UPF complex in the absence of UPF3B, UPF3A also shows increased co-IP with core EJC factor EIF4A3 (Figure 2C) and peripheral protein CASC3 (Figure S2A) in UPF3B knockout cells.

**Figure 2.**
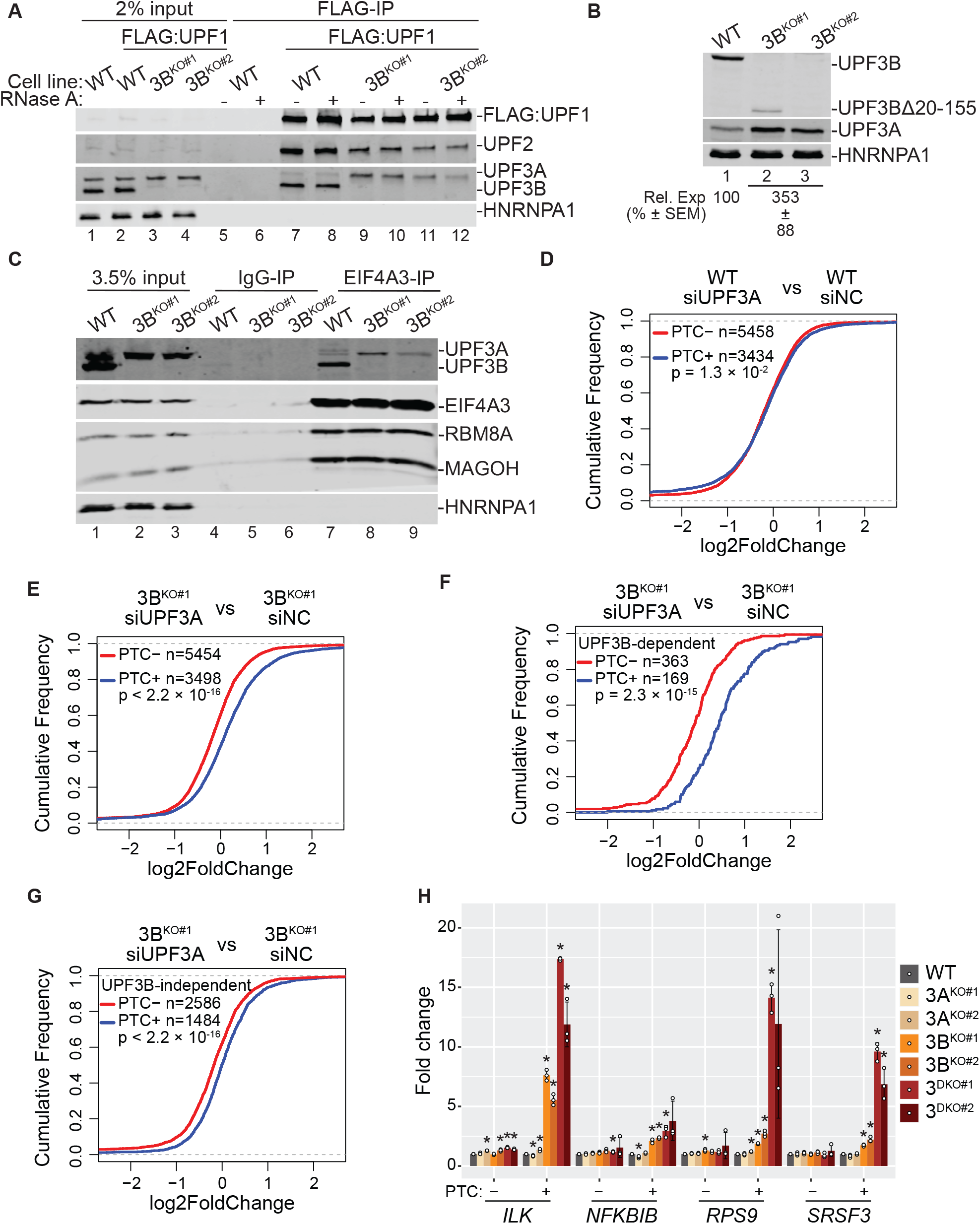
UPF3A activates NMD in the absence of UPF3B. A. Immunoblots showing levels of proteins on the right in input or FLAG immunoprecipitates (FLAG-IP) from WT and UPF3B-KO cells expressing endogenously FLAG-tagged UPF1 protein as indicated above each lane. The presence of RNase A during FLAG-IP is indicated above each lane. B. Immunoblots showing levels of proteins on the right in cells indicated above each lane. At the bottom are relative UPF3A levels after normalization to HNRNPA1 levels (n=4). C. Immunoblots showing levels of proteins (right) in input and immonoprecipitates from IP with normal rabbit IgG (IgG-IP) or antibody targeting EIF4A3 (EIF4A3-IP) from WT and UPF3B-KO cells. D, E. CDF plots of PTC+ isoforms and PTC-isoforms from same set of genes. X-axis represents fold change upon UPF3A knockdown (siUPF3A) versus negative control knockdown (siNC) in, (D) WT cells, and (E) 3B^KO#1^ cells. Number of transcripts in each set (n) and p-value from KS test comparing the two distributions are shown on each plot. F, G. CDF plots of UPF3B-dependent (F) and -independent (G) PTC+ isoforms and PTC-isoforms from same set of genes. X-axis represents fold change upon UPF3A knockdown (siUPF3A) versus negative control knockdown (siNC) in 3B^KO#1^ cells. Number of transcripts in each set (n) and p-value from KS test comparing the two distributions are shown on each plot. H. Bar plot showing average fold change as measured by isoform specific RT-qPCR of PTC+ and PTC-isoform from genes indicated on the bottom in WT and two independent clones of 3A^KO^, 3B^KO^, and 3^DKO^ cells. Relative levels from each replicate are shown by white circles. Error bars indicate standard error of means. The asterisk (*) represents p<0.05 in t-test with null hypothesis of true mean being 1 (n=3).

To validate that UPF3A is indeed incorporated into EJC-UPF complex, we inserted a FLAG affinity tag into the *MAGOH* locus and a MYC affinity tag into the *UPF2* locus. We then knocked out either UPF3A or UPF3B in this dual-tagged cell line by inserting an antibiotic resistance gene followed by polyadenylation signal. From these cells, a tandem IP of FLAG-MAGOH followed by MYC-UPF2 can isolate the EJC-UPF complex from WT, UPF3A knockout or UPF3B knockout cells. In the WT and UPF3A knockout cells, UPF3B is the major paralog incorporated into the EJC-UPF complex as indicated by co-IP of UPF3B and CASC3 (Figure S2B). In comparison, in the UPF3B knockout cells, UPF3A is incorporated into the complex at a much higher level (Figure S2B). We also notice that there is an overall decrease in the abundance of the UPF2-EJC complex in UPF3B knockout cells as compared to WT or UPF3A knockout cells (Figure S2B). Together, these data suggest that UPF3A is capable of simultaneously engaging with the UPF and EJC proteins particularly in the absence of UPF3B.

### UPF3A compensates for UPF3B function in NMD

Previous evidence from human cells suggests that UPF3A can act as a weak NMD activator particularly in cells with reduced UPF3B levels (Chan et al., 2009; Kunz et al., 2006). Interestingly, when overexpressed in wild-type cells, UPF3A can inhibit NMD (Chan et al., 2009; Shum et al., 2016). Our results above suggest that UPF3A may provide UPF3 function during NMD in the absence of UPF3B. To evaluate contribution of UPF3A to NMD in wild-type and UPF3B knockout cells, we knocked-down UPF3A in WT and 3B^KO#1^ cells (Figure S2C) and performed RNA-Seq to quantify global transcript levels as above. We observe that while UPF3A knockdown in WT HCT116 cells leads to widespread changes in the transcriptome (Figure S2D), changes between PTC+ and PTC-isoforms are negligible (Figure 2D). These data suggest that in wild-type HCT116 cells, UPF3A is largely inconsequential for NMD and neither acts as an NMD enhancer or repressor. In contrast to the wild-type cells, when abundance of PTC+ and PTC-transcripts is compared in UPF3B knockout cells after UPF3A or control knockdown, we observe a specific and significant upregulation of the PTC+ transcript group in cells depleted of both UPF3A and UPF3B as compared to cells lacking only UPF3B (Figure 2E). These data suggest that UPF3A possesses the ability to activate NMD that becomes prominent only in the absence of UPF3B while in wild-type cells UPF3A potentially has a function outside NMD.

We next tested if UPF3A and UPF3B affect similar or distinct set of NMD targets. We defined UPF3B-dependent targets as PTC+ transcripts that show significant and ≥1.5-fold upregulation in 3B^KO#1^ cells as compared to WT cells, and compared their change upon additional UPF3A knockdown versus control knockdown in 3B^KO#1^ cells. As control, we compared change in the corresponding PTC-group under the same conditions. We find that UPF3B-dependent PTC+ group shows a strong upregulation after UPF3A knockdown in 3B^KO#1^ cells (Figure 2F). At the same time, the UPF3B-independent transcripts, which change ≤1.2-fold in 3B^KO#1^ cells as compared to WT cells, are also similarly affected by UPF3A knockdown in 3B^KO#1^ cells, albeit to a lesser extent (Figure 2G). Thus, in the absence of UPF3B, UPF3A acts on a similar set of mRNAs as UPF3B, and NMD targets that are insensitive to UPF3B are only weakly affected by UPF3A.

To further validate the UPF3A function in NMD, we created UPF3A knockout cells (3A^KO^) either by deleting the genomic region encompassing exons 1 and 2 and part of intron 2 or by inserting a blasticidin resistance marker followed by a polyadenylation signal (Figure S2E,F). We also created a UPF3A+3B double knockout (3^DKO^) cells by combining the two antibiotic resistance markers for each gene (Figure S2F). qPCR based quantification of PTC+ transcripts that are upregulated upon UPF3B loss (Figure 1E and Figure S1F) and their corresponding PTC-isoforms shows that the loss of UPF3A has minimal or no effect on the abundance of any of the PTC+ isoforms (Figure 2G and S2F) confirming that UPF3A is dispensable for NMD of this subset of transcripts. In comparison, all the PTC+ transcripts show the highest upregulation in the UPF3 double knockout cell lines (Figure 2G and Figure S2F) reflecting the additive effects of loss of UPF3A and UPF3B on NMD of these transcripts (except for *NFKBIB*, which shows a minor additive effect). Together, these results show that UPF3A works in the EJC-dependent NMD but only in the absence of UPF3B. Hereafter, we will use UPF3 to refer to both the paralogs.

We next tested if the minimal effect of the loss of UPF3B on long 3′UTR-containing mRNAs (Figure S1G,H) was due to the compensation of UPF3 function by UPF3A in UPF3B knockout cells. We observe that UPF3A knockdown in UPF3B knockout cells or in WT cells does not lead to an upregulation of long 3′UTR transcripts (Figure S2H). In turn, UPF3A might even mildly inhibit long 3′UTR-mediated NMD as UPF3A knockdown leads to a modest but significant downregulation of the transcript group with the longest 3′UTRs in both WT and UPF3B knockout HCT116 cells (Figure S2I). Due to the minimal impact of UPF3 proteins on long 3′UTR-mediated NMD in HCT116 cells, from hereon we focus only on UPF3 function in EJC-mediated NMD.

### CASC3-containing EJC potentiates UPF3-dependent NMD

We next tested the contribution of other factors from the EJC-mediated NMD pathway to UPF3 function in this branch of NMD. We first focused on CASC3-containing EJC as previous observations from human embryonic kidney (HEK293) cells suggest that there exists a functional synergy between the CASC3-containing EJC and UPF3B (Gerbracht et al., 2020; Mabin et al., 2018). To test if CASC3-UPF3B preferential association is also present in other cell types, we overexpressed in HeLa cells either wild-type CASC3 or a mutant CASC3 (HDAA) that is unable to associate with the EJC core (Ballut et al., 2005). We observe an enhanced EJC-UPF3B association in HeLa cells when wild-type CASC3 but not the CASC3 HDAA mutant is overexpressed (Figure 3A, compare lanes 9-10 with lanes 11-12). To test if CASC3 overexpression can similarly enhance EJC-UPF3A association, we created a UPF3B knockout in HeLa cells by deleting a ∼1.4 kb region of the *UPF3B* locus encompassing part of its promoter region, first exon and part of the first intron (Figure S3A and S3B). UPF3B knockout in HeLa cells results in a similar enhancement of EJC-UPF3A association (Figure S3C) as observed in HCT116 cells (Figure 2C). Furthermore, overexpressing wild-type CASC3 but not the CASC3 HDAA mutant in UPF3B knockout HeLa cells enhances UPF3A co-IP with EIF4A3 (Figure 3B, compare lanes 9-10 with lanes 11-12). Additionally, CASC3 knockdown reduces both EJC-UPF3B and EJC-UPF3A association in wild-type and UPF3B knockout HeLa cells (Figure S3D). Together, these results suggest that CASC3 promotes the EJC-UPF3 association.

**Figure 3.**
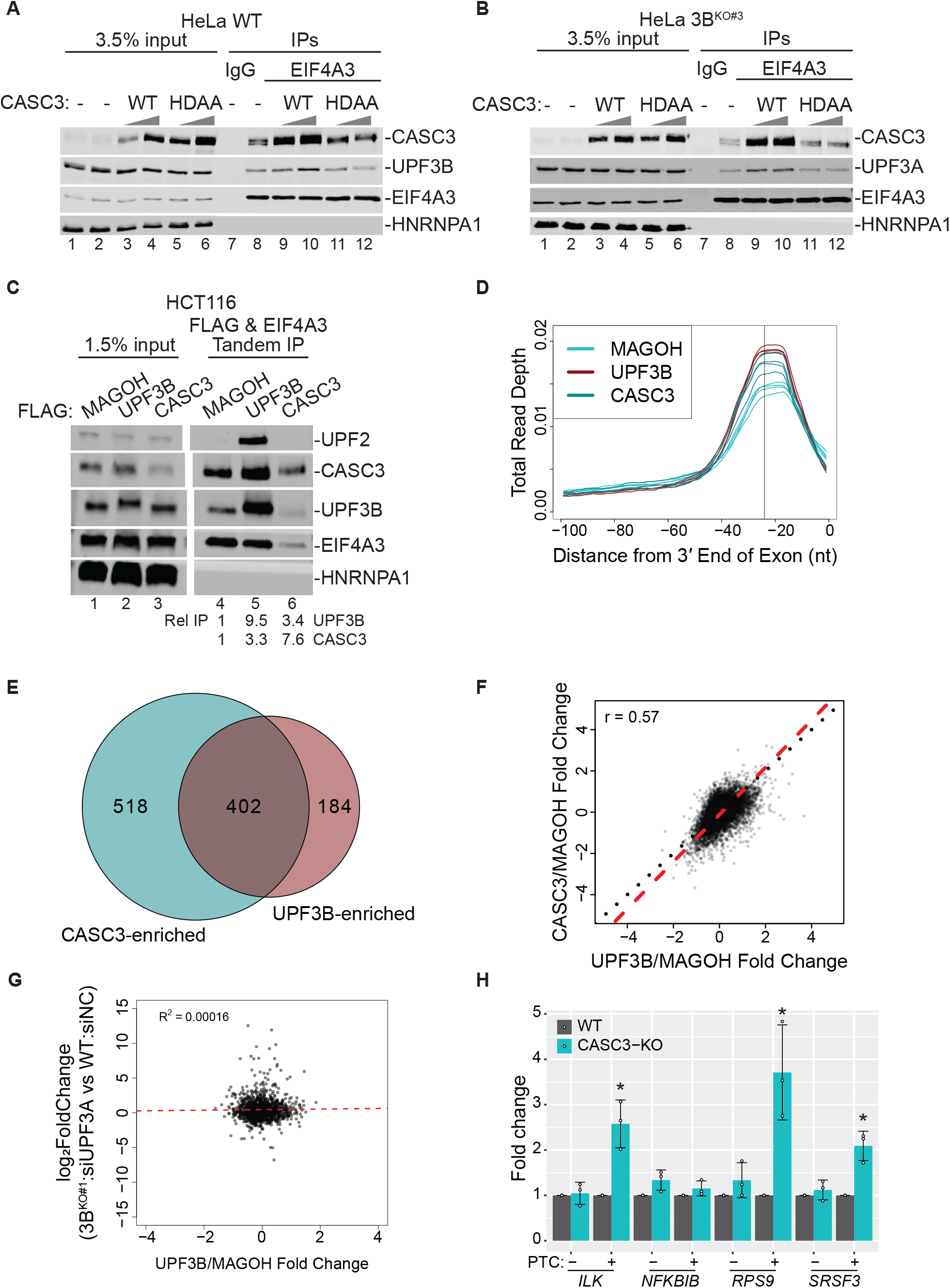
CASC3 contributes to UPF3-dependent NMD. A, B. Western blots showing levels of EJC proteins or HNRNPA1 (control) in input, IgG IP or EIF4A3 IP following overexpression (OE) of CASC3 wild-type (WT) and EJC binding deficient (HDAA) mutant proteins in (A) HeLa Tet-off cells, and (B) 3B^KO^ HeLa Tet-off cells. Ramps above lanes indicate expression levels of the CASC3 proteins. C. Western blots showing levels of EJC/UPF proteins and HNRNPA1 in input and FLAG followed by EIF4A3 tandem IP from HCT116 cells expressing the FLAG-tagged protein indicated above each lane. Quantifications of UPF3B and CASC3 protein enrichment from two replicates are shown at the bottom. D. Meta-exon plot showing read distributions within the 100 nucleotide (nt) window from the exon 3′ end in RIPiT-Seq replicates of MAGOH:EIF4A3, UPF3B:EIF4A3, and CASC3:EIF4A3. The black vertical line indicates the -24 nt position. E. Venn diagram showing the degree of overlap between genes significantly enriched in CASC3:EIF4A3 EJC and UPF3B:EIF4A3 EJC occupancy as compared to MAGOH:EIF4A3 EJC occupancy. F. Scatter plot comparing log2-transformed fold change in occupancy of CASC3:EIF4A3 EJC as compared to MAGOH:EIF4A3 EJC (x-axis) and UPF3B:EIF4A3 EJC compared to MAGOH:EIF4A3 EJC. Each dot represents a gene where gene-level occupancy of each EJC composition was quantified at the canonical position for EJC footprints. Pearson correlation coefficient is shown on the top left. G. Scatter plot showing a comparison between relative UPF3B occupancy on gene (UPF3B:EIF4A3 RIPiT-Seq normalized to MAGOH:EIF4A3 RIPiT-Seq) on x-axis and NMD efficiency of each gene on the y-axis. For each gene in this analysis, NMD efficiency is the highest fold change (in 3B^KO#1^ siUPF3A to WT siNC) observed for its PTC+ isoform. R^2^ from the linear regression fit is shown on the top left. H. Bar plots showing fold changes measured by isoform specific RT-qPCR of PTC+ and PTC-isoform from genes indicated on the bottom in WT and CASC3-KO HCT116 cells. Relative levels from each replicate are shown by white circles. Error bars indicate standard error of means. The asterisk (*) represents p<0.05 in t-test with null hypothesis of true mean being 1 (n=3).

To investigate CASC3-UPF3B association in HCT116 cells, we inserted FLAG sequence into endogenous *UPF3B, CASC3* and *MAGOH* loci to express FLAG-tagged translational fusions of these factors. From these cells, we performed FLAG IPs followed by a second IP of the EJC core factor EIF4A3 to isolate compositionally different EJCs. We find that CASC3 is strongly enriched in UPF3B-containing EJC as compared to EJCs purified via its core factors, which likely are a mixture of EJCs of distinct compositions (Figure 3C, compare lanes 4 and 5). We hypothesized that preferential association between CASC3 and UPF3B will enrich the EJCs containing the two proteins on a similar set of transcripts. To test this possibility, we carried out RIPiT-Seq (RNA IP in tandem followed by high-throughput sequencing (Singh et al., 2012)) to identify the RNA footprints of FLAG-MAGOH:EIF4A3, FLAG-UPF3B:EIF4A3 and FLAG-CASC3:EIF4A3 complexes. As expected, the RNA footprints of the three complexes show a strong enrichment at the exon 3′ ends at the expected EJC binding site (Figure 3D and Figure S3E). To test if UPF3B and CASC3 exhibit synergistic binding to transcripts, we first individually compared CASC3:EIF4A3 and UPF3B:EIF4A3 occupancy to the EJC core (MAGOH:EIF4A3) occupancy. Genes that are enriched in either UPF3B-EJC or CASC3-EJC as compared to the EJC core show a large and significant overlap (Figure 3E). In contrast, little overlap is detected between genes that are UPF3B-depleted and CASC3-enriched, or vice versa (Figure S3F,G). Strikingly, we observe a strong positive correlation between transcriptome wide UPF3B and CASC3 binding relative to the EJC core (Figure 3F) suggesting that UPF3B and CASC3 preferentially bind to a similar set of transcripts.

We next asked if UPF3B-EJC occupancy influences NMD efficiency, i.e., does increased UPF3B-EJC binding to an mRNA lead to more efficient NMD? For this analysis, we selected genes that express at least one PTC+ isoform, and for NMD efficiency of each gene, we used the highest fold change observed for its PTC+ isoform in UPF3 depleted cells (UPF3A knockdown in 3B^KO#1^ cells) as compared to control (WT cells with negative control knockdown). Upon comparison of NMD efficiency estimates for these genes to their EJC core-normalized UPF3B-EJC occupancy, we do not observe any appreciable correlation between the two metrics (Figure 3G). Similar relationship is seen between NMD efficiency and mRNA expression-normalized UPF3B (Figure S3H) or EJC core (Figure S3I) occupancy. Together, these data suggest that in wild-type HCT116 cells UPF3B or EJC occupancy are unlikely to be limiting factors for NMD.

To directly investigate the dependence of UPF3-dependent NMD on CASC3, we measured the levels of UPF3B-dependent PTC+ transcripts in a HCT116 cell line where CASC3 expression is completely knocked out by frameshifting indels around its start codon (Figure S3J). We observe a moderate increase in abundance of *ILK, RPS9* and *SRSF3* PTC+ transcripts while no such change is observed in the case of *NFKBIB* PTC+ isoform, an NMD target that is the least affected by the loss of UPF3 (Figure 2G). Thus, normal CASC3 levels are important for efficient UPF3-dependent NMD.

### UPF2 and UPF3 function in EJC-mediated NMD is interdependent

To test if the UPF3 paralogs are required by UPF2 for its function in EJC-mediated NMD, we knocked down UPF2 in wild-type and in UPF3A and UPF3B double knockout HCT116 cells (Figure S4A). While UPF2 knockdown in wild-type cells alters expression of more than two thousand transcripts (Figure S4B), only less than a quarter are changed after UPF2 knockdown in UPF3 double knockout cells (Figure S4C), suggesting that UPF2 depletion in cells lacking UPF3 causes fewer additional changes in gene expression. Such an effect is more obvious for EJC-mediated NMD substrates. While a significant upregulation of PTC+ isoforms is seen upon UPF2 knockdown in wild-type cells (Figure 4A), no such change is observed between PTC+ and PTC-isoforms upon depleting UPF2 in UPF3 double knockout cells (Figure 4B). These data suggest that UPF2 function in EJC-mediated NMD depends on UPF3, and that these factors act on a shared set of transcripts in the EJC-mediated NMD branch.

**Figure 4.**
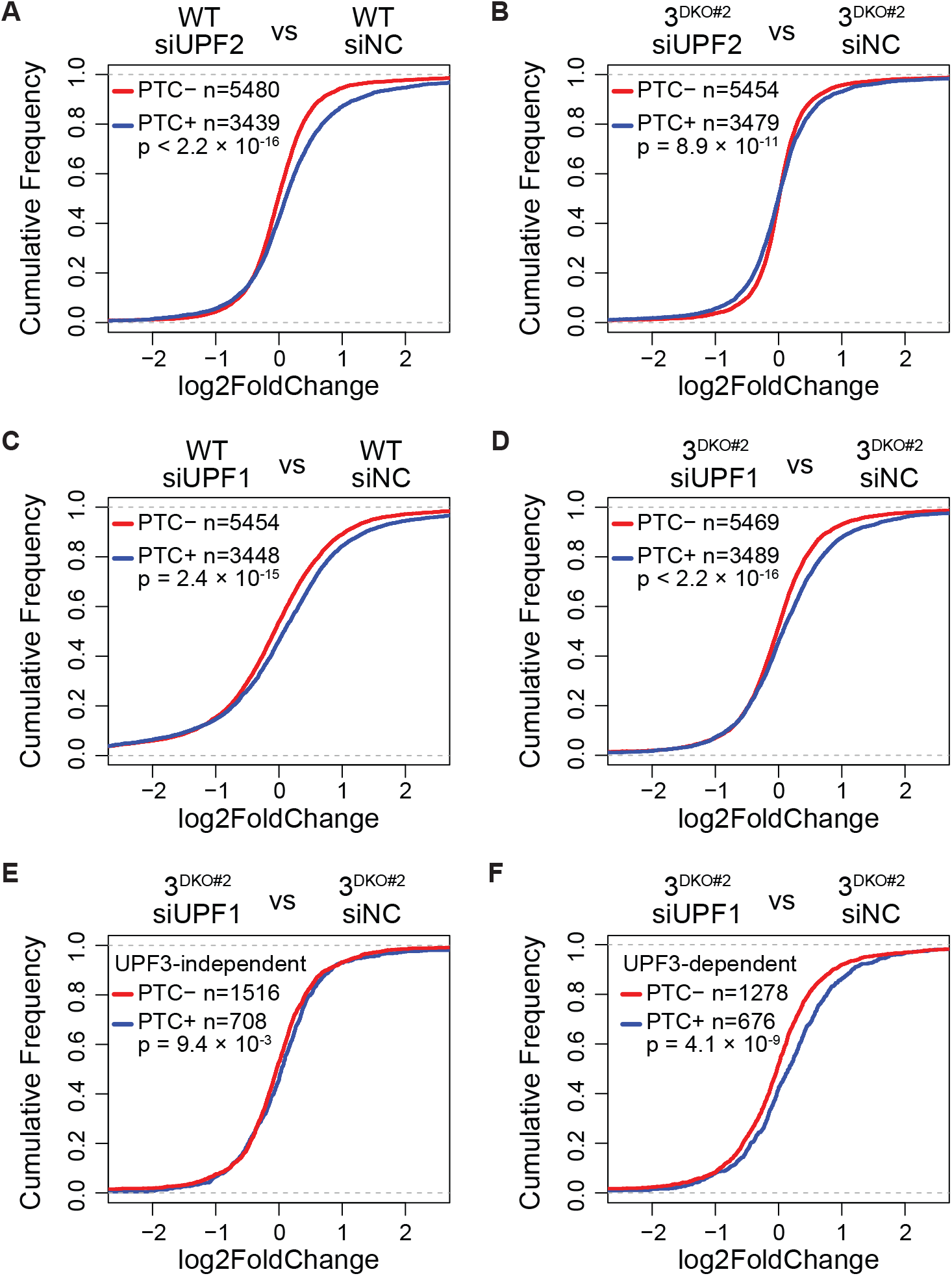
NMD activity in human cells in the absence of both UPF3 paralogs. A, B. CDF plots of PTC+ and PTC-isoforms from same set of genes. X-axis represents log_2_ fold change upon UPF2 knockdown as compared to control knockdown in, (A) WT cells, and (B) 3^DKO#2^ cells. C, D. CDF plots of PTC+ and PTC-isoforms from same set of genes. X-axis represents fold change upon UPF1 knockdown as compared to control knockdown in, (A) WT cells, and (B) 3^DKO#2^ cells. (Figure 4A is the same as Figure 1D.) E, F. CDF plots of UPF3B-independent (E) and -dependent (F) PTC+ and PTC-isoforms from same set of genes. X-axis represents fold change upon UPF1 knockdown (siUPF1) versus negative control knockdown (siNC) in 3^DKO#2^ cells. Number of transcripts in each set (n) and p-value from KS test comparing the two distributions are shown on each plot.

### EJC-mediated NMD can completely bypass the need for UPF3

It has been previously reported that NMD of a TCR-β reporter mRNA and a handful of endogenous NMD substrates can occur independently of UPF3 in human cells (Chan et al., 2007). However, in these experiments UPF3 paralogs were depleted using RNA interference thus leaving open a possibility that some residual UPF3 proteins may still be able to sustain NMD. We therefore tested the existence and extent of any EJC-mediated NMD that can still occur in UPF3 double knockout cells. We observe that, like in wild-type cells (Figure 4C), UPF1 knockdown in UPF3 double knockdown cells leads to further upregulation of PTC+ isoforms compared to PTC-isoforms (Figure 4D), suggesting that some EJC-mediated NMD can still function in the complete absence of both UPF3 proteins. Our data provides a strong support for the existence of a UPF3-independent NMD branch, which also is unlikely to require UPF2.

It remains unknown if UPF3-dependent and -independent NMD branches target different mRNAs or if the two branches target same set of mRNAs that show variable NMD commitment in the presence/absence/variable levels of UPF3 proteins. To test this idea, we separated PTC+ mRNAs into two groups (i) a UPF3-dependent group that is significantly upregulated ≥1.5-fold in 3^DKO^ cells, and (ii) a UPF3-independent group that changes ≤1.2-fold in 3^DKO^ cells. If UPF3-independent NMD branch targets distinct mRNAs that are still degraded by NMD in 3^DKO^ cells, then these PTC+ mRNAs are expected to be upregulated upon UPF1 knockdown in 3^DKO^ cells. However, we observe that the UPF3-independent PTC+ mRNAs show only a very minor upregulation as compared to their PTC-counterparts when 3^DKO^ cells are depleted of UPF1 (Figure 4E). In contrast, the UPF3-dependent group of PTC+ mRNAs shows a more prominent upregulation under these conditions (Figure 4F). These data indicate that PTC+ mRNAs that undergo UPF3-dependent NMD can still be targeted by the NMD pathway in UPF3-independent manner, perhaps at a reduced rate.

### UPF3A and UPF3B differ in their NMD activity that is dictated by their mid domains

Although previous work suggests that in human cells UPF3A can suppress NMD of certain endogenous genes (Shum et al., 2016), we did not observe such an activity of UPF3A in our analysis of EJC-mediated NMD targets in HCT116 cells (Figures 2 and S2). Notably, among previous reports of UPF3A’s NMD suppressing activity in human cells, most robust NMD inhibition was observed in the case of a β-globin NMD reporter mRNA under UPF3A overexpression conditions (Chan et al., 2009; Shum et al., 2016). Indeed, we confirmed that β-globin mRNA reporter with a PTC at codon 39 (β39) is stabilized when UPF3A is overexpressed in wild-type HeLa cells. In comparison, UPF3B overexpression shows little effect on the reporter mRNA half-life as compared to the control (Figure 5A). Additionally, these results also highlight that despite UPF3A’s ability to compensate for UPF3B function in NMD, the two paralogs have notable differences in their NMD activity. To further investigate these differences, we compared the ability of UPF3A and UPF3B to rescue the strong defect in β39 reporter mRNA NMD in UPF3B knockout HeLa cells as compared to the wild-type cells (Figure 5B). While overexpressing exogenous UPF3B in UPF3B knockout cells fully rescues β39 reporter mRNA decay, overexpressing UPF3A in these cells only mildly rescues the decay of the reporter (Figure 5B). Together, these data suggest that, while UPF3A can functionally compensate for UPF3B, the two paralogs exhibit different NMD activities.

**Figure 5.**
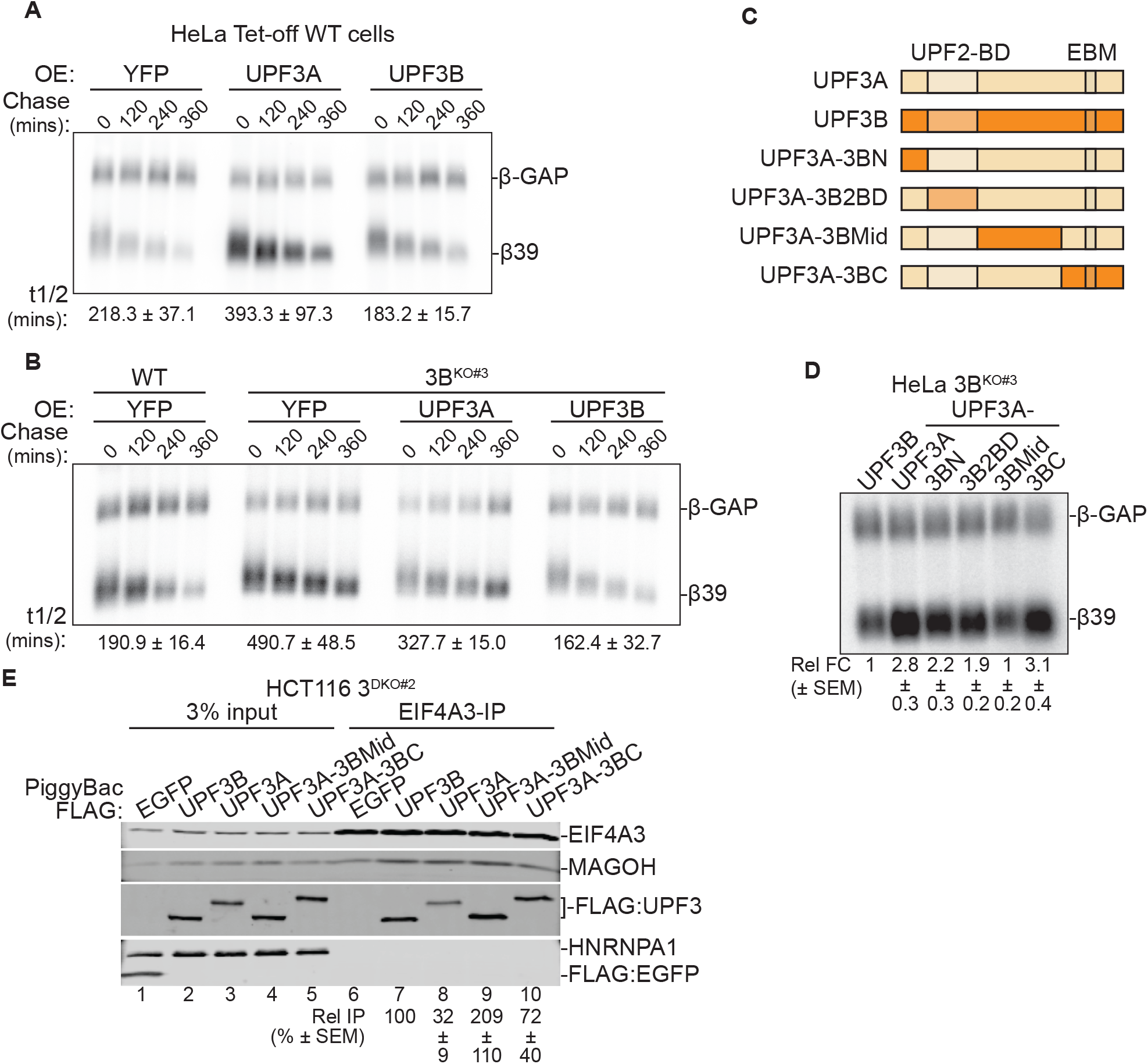
Human UPF3 paralogs differ in NMD activity. A, B. Northern blots showing levels of β-globin reporter mRNAs in, (A) wild-type HeLa Tet-off cells, and (B) UPF3B knockout HeLa Tet-off cells. β39 is a tetracycline (Tet)-inducible reporter with a PTC at codon 39 whose levels are shown at different timepoints after transcriptional shut-off (chase) as indicated above each lane. β-GAP is a stable, constitutively-expressed, longer β-globin mRNA used as transfection control. Proteins overexpressed (OE) in each condition are indicated on top and reporter mRNA half-lives (t_1/2_) along with standard error of means are on the bottom. C. Schematic of human UPF3A, UPF3B and the UPF3A chimeric proteins where UPF3A domains are replaced by the corresponding domains from UPF3B (see material and methods for detailed domain definition). Previously characterized UPF2 binding domain (UPF2-BD) and EJC-binding motif (EBM) are shown. D. Northern blot showing steady-state levels of β39 NMD reporter and β-GAP control in HeLa Tet-off UPF3B knockout cells upon overexpression of wild-type UPF3 proteins or different UPF3A chimeric proteins indicated above each lane. Below each lane, relative fold-change (Rel. F.C.) indicates β39 reporter levels (normalized to β-GAP control) as compared to the normalized β39 reporter levels in UPF3B expressing cells. E. Immunoblot showing levels of EJC proteins or HNRNPA1 in input or EIF4A3-IP from 3^DKO#2^ cells expressing different UPF3 proteins or EGFP as a control as indicated above each lane. Relative IP of FLAG-tagged proteins are quantified against EIF4A3.

We next sought to identify the molecular basis of the differences in the NMD activity of the two human UPF3 paralogs. We created a series of domain swap mutants where each UPF3A domain is replaced by the corresponding sequence from UPF3B (Figure 5C) with the goal to identify the UPF3B domain that will confer UPF3A with a full UPF3B-like NMD activity. An expectation based on the previous work is that the lower UPF3A NMD activity results from its weaker EJC binding compared to UPF3B due to an arginine-to-alanine change at position 423 in the EJC binding motif (EBM) within the carboxy (C)-terminal domain (Kunz et al., 2006). However, we find that in the UPF3B knockout cells overexpressing a UPF3A mutant with the UPF3B C-terminal domain, the steady-state β39 mRNA levels are similar to those in the cells transfected with wild-type UPF3A and ∼3-fold higher than the cells expressing UPF3B (Figure 5D). Surprisingly, the UPF3A mutant that carries the UPF3B mid-domain (region between the UPF2 binding domain and the C-terminal domain) lowers the β39 mRNA steady-state levels to a similar extent as UPF3B (Figure 5D). Thus, we conclude that the UPF3B mid-domain, and not its EBM-containing C-terminal domain, might underlie the difference between UPF3A and UPF3B NMD activity.

To compare the EJC and UPF binding ability of the UPF3A swap mutants to the wild-type UPF3 paralogs, we created stable cell lines in the UPF3 double knockout background using the PiggyBac transposon system to express FLAG-tagged UPF3 proteins and the UPF3A swap mutants. EIF4A3 IP from these cells shows that, as previously reported, UPF3A shows weaker binding to the EJC as compared to UPF3B (Figure 5E, compare lanes 7 and 8). Interestingly, both UPF3A-3BMid and UPF3A-3BC mutants show an increased association with the EJC (Figure 5E and Figure S5A) even though only UPF3A-3BMid mutant can rescue the decay of NMD reporter mRNA (Figure 5D). We conclude that while the UPF3B mid and C-terminal domains can independently enhance EJC association, possibly via distinct mechanisms, the difference in NMD activity of UPF3A and UPF3B primarily stems from their mid-domains. A previous report has suggested that UPF3B mid-domain (smaller than as defined here) can associate directly with eRF3 (GSPT1) protein in vitro (Neu-Yilik et al., 2017). However, FLAG-IP of UPF3A or UPF3B fails to co-IP detectable eRF3 (Figure S5B), possibly due to the transient nature of such association in HCT116 cells.

### EJC binding is largely dispensable for UPF3 NMD activity

While we do observe stabilization of NMD mRNA after UPF3A overexpression, our data cannot conclude if such “NMD repressor” activity is physiologically present. Interestingly, sequence alignment of human, mouse and rat UPF3 C-terminal domains reveals that while the domain is conserved among the three species, mouse and rat UPF3A lack most or all residues required for EJC-binding whereas human UPF3A retains most of the EJC-binding residues (Figure 6A). Consistently, IP of FLAG-tagged mouse UPF3A (mUPF3A) from 3^DKO^ cells shows a near complete absence of EJC factors in the immunoprecipitates, while human UPF3A (hUPF3A) can still associate with EJC proteins albeit more weakly as compared to human UPF3B (Figure 6B). Importantly, all three proteins show a comparable association with UPF2. Furthermore, mUPF3A transiently expressed in HeLa cells also fails to co-IP any detectable levels of EJC proteins (Figure S6). We conclude that over the course of evolution, the UPF3A proteins in mouse, and most likely in rat as well, have lost their EJC binding ability.

**Figure 6.**
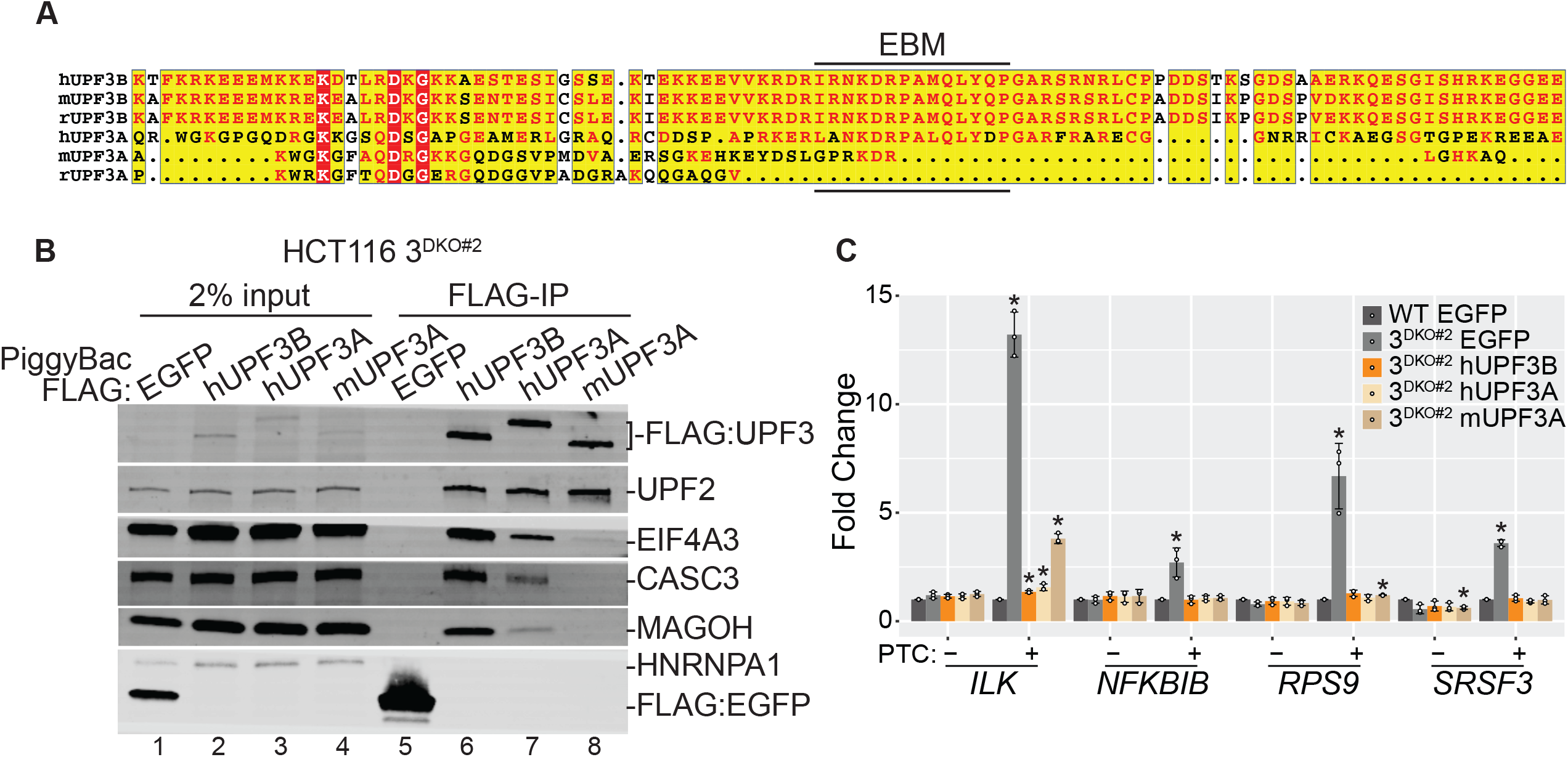
EJC binding is dispensable for NMD activity of UPF3. A. Protein sequence alignment of UPF3 C-terminal regions from different mammalian species. B. Immunoblot showing levels of EJC and UPF proteins (on the right) in input or FLAG-IP samples from 3^DKO#2^ cells expressing different FLAG-tagged proteins indicated above each lane. C. Bar plots showing isoform specific RT-qPCR-based measurement of relative levels of PTC+ and PTC-isoforms of genes indicated below from wild-type (WT) or 3^DKO#2^ cells expressing the specified proteins. Relative levels from each replicate are shown by white circles. Error bars indicate standard error of means. The asterisk (*) represents p<0.05 in t-test with null hypothesis of true mean being 1 (n=3).

We hypothesized that due to its loss of EJC binding activity, mouse UPF3A may not be able to compensate for *UPF3B* loss in human cells, which might reconcile the difference between our observation and the previous study (Shum et al., 2016). Surprisingly, however, when we expressed mouse or human UPF3A proteins, or human UPF3B as a control, in 3^DKO^ cell lines, all three UPF3 proteins fully rescue the NMD of all but one PTC+ isoforms we examined (Figure 6C). Mouse UPF3A expression in 3^DKO^ cells leads to partial rescue only in the case of the *ILK* PTC+ isoform, which shows the strongest UPF3B dependence (Figures 1E, 2G and 6C) and full rescue by either human UPF3A or UPF3B (Figure 6C). These data provide the first evidence that EJC-binding is not required for the NMD activity of UPF3 in human cells and may play a more secondary role in EJC-mediated NMD.

## DISCUSSION

Of the three core NMD factors, UPF3 has evolved most rapidly in eukaryotes. In multicellular organisms, on the one hand it appears to have lost its essentiality for NMD activity, and on the other, it has gained an ability to interact with the NMD-stimulating EJC. Furthermore, in vertebrates, *UPF3* gene has duplicated into paralogous *UPF3A* and *UPF3B*, which have diverged in their EJC binding ability. The emergence of these variations in *UPF3* raises several questions: What is UPF3’s primary function in NMD? If NMD can occur without EJC in yeast, what is the role of EJC interaction in UPF3 function? How do the two paralogs contribute to UPF3 activity in the pathway? How can NMD function in the absence of UPF3B, or UPF3 altogether, and how prevalent is such an NMD activity? Our work here using complete UPF3A and UPF3B loss-of-function human cell lines adds to the experimental evidence that is leading toward answers to these important questions. Based on the existing data and our results reported here, we present an updated model of UPF3A and UPF3B function in mammalian NMD (Figure 7), which is further elaborated below.

**Figure 7.**
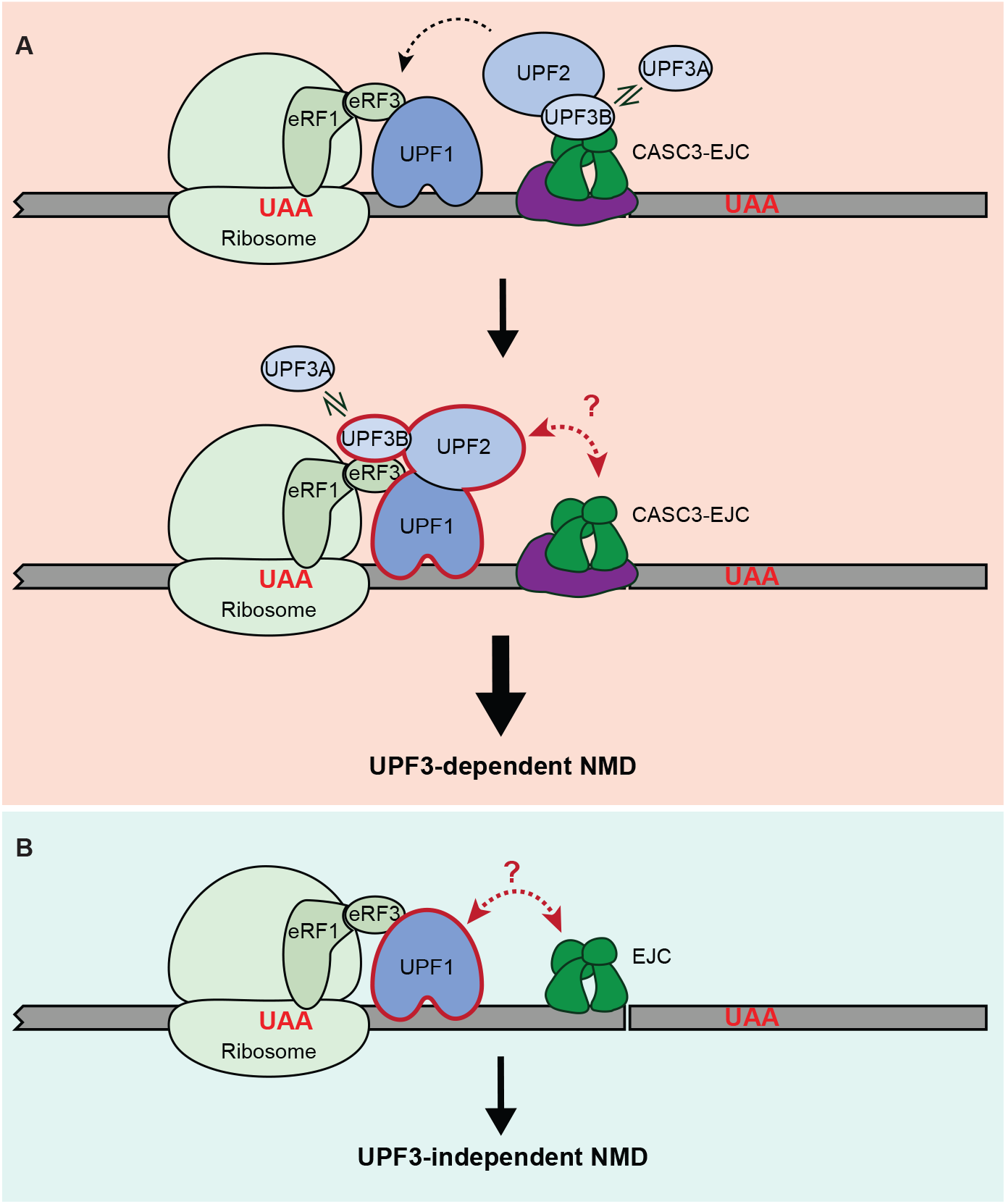
A model for UPF3 function in EJC-enhanced NMD. A. Top: In UPF3-dependent NMD, prior to UPF1 activation, CASC3-EJC enhances the presence of UPF3B and UPF2 on exon-exon junctions. UPF3A can replace UPF3B when UPF3B levels are insufficient. Such enhanced concentration of UPF3 and UPF2 in 3′UTR can later facilitate the formation of NMD complex with UPF1 (dashed arrow at the top). Bottom: During NMD activation, UPF3-eRF3 association is likely to play an important role in sensing aberrant translation termination. While EJC might still play a role during NMD activation, its association with UPF3 is dispensable for NMD. Red double-headed arrow signifies possible UPF complex-EJC communication independently of UPF3. B. NMD can occur in UPF3-independent manner. It remains possible that EJC can still communicate with premature termination complex in a UPF3-, and UPF2-independent manner to elicit mRNA decay (red dashed double-headed arrow).

### UPF3A is an NMD activator

The early studies on UPF3A suggested that it acts as a weak NMD activator in human cells (Chan et al., 2009; Kunz et al., 2006). However, the recent work by Shum et al using mouse models and cell lines has instead suggested that the UPF3 gene duplication fueled neo-functionalization of UPF3A into an NMD repressor (Shum et al., 2016). In the model proposed by Shum et al, weak EJC binding by UPF3A sequesters UPF2 away from the NMD complex thereby leading to NMD inhibition (Shum et al., 2016). In our work here in human (HCT116) cells, we do not observe any negative (or positive) effects on the levels of PTC-containing mRNAs when UPF3A is depleted via RNA interference (Figure 2D) or when it is completely knocked out (Figure 2G) in wild-type cells. These data suggest that UPF3A does not interfere with UPF3B function in EJC-mediated NMD, and hence does not act as NMD repressor in these cells. On the contrary, several lines of evidence from our work suggests that UPF3A acts as an NMD activator, particularly in the absence of UPF3B. In cells lacking UPF3B, UPF3A is upregulated (Figure 2B) (Chan et al., 2009; Nguyen et al., 2012; Tarpey et al., 2007), and its incorporation in EJC-UPF complexes is dramatically enhanced (Figures 2A,2C, S2A-B) (Chan et al., 2009). Further, a partial or complete UPF3A depletion in UPF3B lacking cells leads to robust upregulation of EJC-mediated NMD targets, both at a global level (Figure 2E) and at an individual transcript level (Figures 2G, S2F). This evidence suggests that in the absence of UPF3B, UPF3A engages with the NMD machinery to sustain the EJC-mediated NMD pathway. Moreover, we find that UPF3A is comparable to UPF3B in its ability to rescue EJC-mediated NMD of various endogenous PTC-containing mRNAs (Figure 6C) although some notable differences between the two paralogs are observed (see below). Additionally, UPF3A’s impact on NMD overlaps with that of UPF3B (Figure 2F) (Figure 2G). Importantly, a parallel study by Wallmeroth et al. also shows that UPF3A functions as a NMD activator in HEK293 cells that lack UPF3B. The redundancy between UPF3A and UPF3B is more exacerbated in these cells as only depletion of both the proteins leads to transcriptome-wide NMD inhibition (Wallmeroth et al., 2021). This evidence also suggests that in human patients with UPF3B inactivating mutations, UPF3A can likely fill in for UPF3B in most of the EJC-dependent NMD, which is critical for several physiological processes (e.g., hematopoiesis (Weischenfeldt et al., 2008)). However, UPF3A cannot compensate for UPF3B functions in select contexts such as brain development possibly due to differences in function or gene expression patterns of the paralogs.

While our findings point to an NMD activating role for UPF3A, in certain specific contexts, UPF3A can inhibit NMD. For example, under artificial conditions, UPF3A overexpression in wild-type HeLa cells slows down NMD of the β-globin reporter mRNA (Figure 5A) (Chan et al., 2009). Such conditions do arise in specific cell types (e.g. mouse germ cells) and/or developmental stages (e.g. early mouse embryogenesis, spermatogenesis) where UPF3A expression is dramatically increased. An important context is male germ cells where UPF3A is likely the main source of UPF3 activity due to the presumed silencing of *UPF3B* due to meiotic X-chromosome inactivation (Turner, 2007). Shum et al showed that high UPF3A to UPF3B ratio in these cells inhibits NMD of selected NMD targets. (Shum et al., 2016). It will be important to assess UPF3A function in NMD in germ cells at a fuller global scale to determine if NMD repression is its dominant function in these cells or if it acts both as NMD activator and repressor, perhaps in transcript specific manner. Notably, Shum et al. did find that UPF3A can activate NMD of a small subset of transcripts in a mouse stem cell line. Our findings here also shed light on the mechanistic basis of potential NMD inhibitory action of UPF3A. We show that mouse UPF3A, which completely lacks EJC binding activity (Figure 6B), can still rescue NMD of most endogenous mRNA targets in UPF3 double knockout cells to the same level as the human UPF3A or UPF3B proteins. This ability of mouse UPF3A to activate NMD without EJC binding suggests that it is unlikely that weak or no EJC binding by UPF3A proteins can cause NMD repression. Instead, UPF3A may inhibit NMD potentially by outcompeting the stronger NMD factor UPF3B (Figure 5A).

The basis of conservation of two similar UPF3 paralogs remains to be fully appreciated. It is possible that in addition to UPF3A’s NMD repressive activity, its ability to activate NMD, or even its NMD-independent functions (Figure S2D; (Ma et al., 2019)) may have contributed to its conservation through vertebrate evolution.

### Dispensability of UPF3-EJC interaction warrants a revised model for UPF3 function in NMD

Current NMD models suggest that in the EJC-mediated NMD, UPF3B acts as a bridging molecule between the UPF proteins and the downstream EJC. This view is based on UPF3B’s ability to directly bind to UPF2 via its N-terminal RRM domain (Kadlec et al., 2004) and to a composite surface on the EJC core via the EJC binding motif in its C-terminus (Chamieh et al., 2008). Surprisingly, we find that mouse UPF3A, which is missing most of the EJC binding motif (Figure 6A) and hence lacks any detectable EJC binding (Figure 6B), is still capable of rescuing NMD of several PTC-containing mRNAs (Figure 6C). Additionally, replacing the weaker EJC binding C-terminal region of human UPF3A with the stronger EJC binding C-terminal domain of human UPF3B does not improve the NMD function of the chimeric UPF3A protein (Figures 5C-E). Thus, our findings suggest that UPF3 proteins (UPF3A or UPF3B) can activate NMD without EJC binding, thereby challenging the decades-old bridging model for UPF3 function in the pathway.

We propose that EJC binding by UPF3 proteins is not a primary activity of these proteins in the NMD pathway. It is more likely that EJC binding is important to recruit UPF3 (and perhaps UPF2) to mRNA exon-exon junctions (Figure 3, Figure 7) to increase the likelihood of NMD activation by yet another UPF3 function when translation terminates at PTC (Figure 7). The recruitment of both human UPF3 paralogs to mRNA via the EJC can be enhanced by CASC3 (Figures 3, S3 and 7), which is a defining component of a compositionally distinct EJC (Mabin et al., 2018). How CASC3 enhances EJC interaction with UPF3 remains to be seen. The current understanding of CASC3 and UPF3B interaction with the EJC core is limited to the small regions of the two proteins that directly contact the EJC (Buchwald et al., 2010; Melero et al., 2012). It is possible that additional interactions between other regions of CASC3 (e.g. N- or C-terminal domains on either side of the EJC binding SELOR domain) and UPF3 (e.g. mid-domain, which enhances EJC interaction (Figure 5D)) contribute to EJC association of UPF3. Although the effect of CASC3 loss on UPF3-mediated NMD targets appear to be modest (Figure 3H), modulation of CASC3 levels, for example by miR128 in neuronal cells (Bruno et al., 2011), can regulate UPF3-dependent NMD. How can the need for UPF3 recruitment to mRNAs via EJC be bypassed? It is possible that at higher UPF3 expression levels, such as those achieved in the rescue experiments in Figure 6, this prior mRNA recruitment of UPF3 becomes dispensable, and the primary function of UPF3 is sufficient to drive NMD. Interestingly, compromised EJC recruitment of UPF3B in cells depleted of ICE1, which also aids in UPF3B-EJC interaction, can be similarly overcome by UPF3B overexpression (Baird et al., 2018).

What is the primary role of UPF3 proteins in NMD activation? Early studies from yeast revealed that these proteins, including UPF3, can physically engage with termination factor eRF3 (Wang et al., 2001). These data suggested a role for UPF proteins in discrimination between normal and premature termination events, but the detail of such mechanisms has remained elusive. A recent investigation using an in vitro assay to monitor translation termination found that UPF3B, but not UPF1 or UPF2, can slow down the termination reaction and promote disassembly of the terminated ribosome (Neu-Yilik et al., 2017). Interestingly, this report shows that UPF3B can directly interact with eRF3 and eRF1. In fact, the direct in vitro UPF3B-eRF3 interaction is mediated by a UPF3B region that falls within the segment that we define here as the mid-domain (Figure 5C). How these translation termination-linked UPF3 activities and interactions contribute to NMD was however unknown. Our work here shows the importance of the mid-domain for efficient NMD (Figure 5D), even though we do not detect an association between UPF3B and eRF3 in HCT116 cells (Figure S5B). It is possible that this interaction is transient in nature and is reliably detectable only when the two proteins are exogenously expressed at much higher levels as in the previous work (Neu-Yilik et al., 2017). It also remains to be seen if differences in NMD activity of UPF3A and UPF3B is governed by the differences in their mid-domains to engage with termination factors. Nonetheless, our data and the findings of Wallmeroth et al. in the accompanying paper along with the published work (Neu-Yilik et al., 2017) indicate that the UPF3 mid-domain plays an important role in NMD activation (Wallmeroth et al., 2021). The functional relevance of this poorly characterized region of the UPF3 proteins is further underscored by several missense *UPF3B* mutations that fall within this domain in individuals with neurodevelopmental disorders (Alrahbeni et al., 2015).

Once translation terminates in a context where signals promoting normal termination are diminished (e.g. weakened PABPC1-eRF interaction, (Ivanov et al., 2008; Peixeiro et al., 2012; Singh et al., 2008)) or where signals promoting premature termination are increased (e.g. more frequent 3′UTR binding of UPF1 and/or increased local concentration of UPF3/UPF2 by the downstream EJC), early steps in the NMD pathway ensue. Perhaps, these include recruitment of UPF3B to the termination complex via its interactions with eRFs, which leads to slowing down of the termination reaction. It is notable that premature termination has been proposed to be more inefficient than normal termination (He & Jacobson, 2015) even though the direct observation and mechanistic basis of such phenomenon remains elusive (Karousis et al., 2020). UPF1 and UPF2 have also been shown to interact with termination factors and may play a role in this process (Ivanov et al., 2008; Kashima et al., 2006; López-Perrote et al., 2016; Singh et al., 2012; Wang et al., 2001). Once termination reaction is deemed to be premature, yet another function of UPF3B can be to promote ribosome dissociation and/or recycling (Neu-Yilik et al., 2017). While it remains to be seen when in relation to termination reaction do critical NMD events such as UPF1 phosphorylation and UPF1 ATPase activation occur, it is likely that UPF3B continues to function at these downstream steps in the NMD pathway. It will be an important goal for future studies to precisely define the order and mechanism of steps that lead to NMD activation, and the contribution of UPF3 proteins to these steps. It will be also important to determine UPF3-independent mechanisms whereby the 3′UTR bound EJCs enhance premature termination and NMD (Figure 7).

### UPF3-dependent and -independent NMD branches

Even though UPF3 (UPF3B) plays a critical role in activating efficient NMD, multiple lines of evidence suggests that NMD in many organisms can proceed independently of UPF3. For example, unlike UPF1 and UPF2, UPF3 is not essential for viability in flies and its loss has only modest effect on many NMD targets (Avery et al., 2011). Similarly, some mRNAs can undergo NMD in human cells with dramatically reduced UPF3B levels or no UPF3B at all (Chan et al., 2007; Nguyen et al., 2012; Tarpey et al., 2007). Moreover, few mRNAs were only mildly affected upon severe depletion of both UPF3A and UPF3B from human cells (Chan et al., 2007, 2009). Our work now provides most definitive evidence for existence for UPF3-independent NMD as the PTC-containing mRNAs can be further upregulated upon UPF1 knockdown in cells completely lacking both UPF3A and UPF3B (Figure 4D). This UPF3-independent NMD is likely to be also UPF2-independent as the same set of transcripts are not affected by UPF2 depletion in UPF3 double knockout cells (Figure 4B). How could NMD proceed in the absence of UPF3 (and UPF2) remains to be investigated.

Intriguingly, we observe that the same set of NMD-targeted transcripts that are UPF3 sensitive can be further upregulated upon UPF1 depletion in UPF3 lacking cells (Figure 4F). Thus, these mRNAs can still undergo EJC-mediated NMD in the absence of UPF3. However, the NMD targets that are unaffected in UPF3 knockout cells remain largely unperturbed as a group under UPF1 limiting conditions (Figure 4E). This suggests that rather than targeting different sets of mRNAs, UPF3-dependent and UPF3-independent NMD branches are more likely to reflect fractions of same mRNAs that commit to NMD in UPF3-dependent or -independent manner. Further, in the parallel study, Wallmeroth et al. show that even UPF3B activities like UPF2 or EJC binding can function in a redundant manner to induce NMD (Wallmeroth et al., 2021). The mechanistic basis and functional significance of such a stochastic/conditional function of NMD factors in the pathway represents yet another exciting frontier for future work.

## MATERIALS AND METHODS

### Plasmids

For CRISPR-Cas9-mediated antibiotic resistance marker and polyadenylation signal knock-in experiments (for gene knockouts), PX330 plasmid was used for introducing Cas9-mediated cuts. PX330 was a gift from Feng Zhang (Addgene plasmid # 42230; http://n2t.net/addgene:42230; RRID:Addgene_42230). Guide RNA sequence was cloned as previously described (Ran et al., 2013). For donor plasmids, 300-500 bp left and right homology region from gene-of-interest, and puromycin resistance marker-Bovine Growth Harmone polyA-signal (amplified from pMK232 (Natsume et al., 2016)) or blasticidin resistance marker-Simian Virus 40 polyA-signal (amplified from pcDNA6/TR (Thermo Fisher)) were cloned into pTwist-Amp (Twist Bioscience) using Golden Gate Assembly (NEB). pMK232 was a gift from Masato Kanemaki (Addgene plasmid # 72834; http://n2t.net/addgene:72834; RRID:Addgene_72834)

For the UPF3 WT and chimeric protein plasmids, human UPF3A (CCDS9543.1) domains, N-terminus (2-62), UPF2-binding domain (63-160), Mid (163-385), and C-terminus (386-476), are replaced by corresponding human UPF3B (CCDS14587.1) domains, N-terminus (2-45), UPF2-binding domain (46-143), Mid (146-370), and C-terminus (371-470). UPF3 and chimeric proteins DNA are cloned into pcDNA3ez-FLAG plasmid using BamHI and XbaI as previously described (Mabin et al., 2018; Singh et al., 2007). For CASC3 expression plasmid, full length CASC3 and CASC3-HDAA mutant are cloned into pcDNA3ez with EcoRI and XbaI.

For PiggyBac transposase expression plasmid, hyPBase sequence (Yusa et al., 2011) was codon optimized for human cell expression and a synthetic DNA was cloned into mammalian expression vector pTwist-CMV-Beta-Globin by Twist Bioscience. PiggyBac transposon plasmids with Tet-ON system are made from PB-TRE-EGFP-EF1a-rtTA plasmid in Addgene #104454, which was a gift from Volker Busskamp (Addgene plasmid # 104454; http://n2t.net/addgene:104454; RRID:Addgene_104454). EGFP inserts in this plasmid were replaced by restriction sites NheI and NotI, and the puromycin resistance marker was replaced by neomycin resistance marker using Gibson Assembly (NEB). FLAG-tagged gene of interests are moved to the PiggyBac plasmid from pcDNA3 using NheI and NotI site.

Plasmids expressing tet-inducible and control β-globin NMD reporters were previously described (Lykke-Andersen et al., 2000; Singh et al., 2007).

### Cell Culture

HCT116 (ATCC) and HeLa Tet-Off (Takara) cell lines were cultured at 37°C and 5% carbon dioxide in a humidified chamber. McCoy’s 5A (Modified) Medium (Gibco) for HCT116 cells and Dulbecco’s Modified Eagle Medium with High Glucose (Gibco) for HeLa Tet-Off cells were supplemented with 10% Fetal Bovine Serum (Sigma) and 1% Penicillin-Streptomycin (Fisher).

### Cell transfection for transient or stable expression

For protein knockdown using siRNA, 1.6 μl of RNAiMAX, 60 pmol of siRNA and 200 μl OMEM was incubated for 20 mins following manufacturer protocol. 3 × 10^5^ cells were then added to the transfection mixture in a 6-well plate. 48 hours after initial transfection, total RNA was harvested.

For transient expression of proteins in HeLa cells, plasmids were transfected using JetPrime (PolyPlus) transfection reagent following manufacturer protocol with one fifth of recommended DNA (e.g. 200 ng DNA was used per well of a 12-well plate if the user manual recommend 1000ng DNA). 24-48 hours later, cells were harvested for immunoprecipitation or northern blot.

To make stable PiggyBac cell lines, 286 ng of pTwist-CMV-BetaGlobin-hyPBase plasmid was co-transfected with 714 ng of transposon plasmid (neomycin resistant) that carries the gene of interest into a 6 well plate with 3 × 10^5^ cells seeded a day before. 48 hours post transfection, cells were trypsinized and expanded under 600 μg/ml G418 selection for 2 weeks. Polyclonal cells resistant to G418 were then expanded and frozen for further experiments. To induce the protein expression from polyclonal PiggyBac stable cells, 100 ng/ml of final doxycycline was added to the medium and cells were harvested after 24 hours for immunoprecipitation or RNA extraction.

### Electroporation

For electroporating CRISPR-Cas9 complexes into HeLa or HCT116 cells, ∼2.5 × 10^5^ cells were washed in PBS and resuspended in Ingenio Electroporation Solution (Mirus) with 1-2 μM RNP complex to a final volume of 50 μl. The electroporation mix was then transferred to a Gene Pulser Electroporation Cuvette (0.2-cm gap). Electroporation was performed with Gene Pulser Xcell Electroporation Systems (Bio-Rad) under the following conditions: HCT 116 cells: 120 V, 13 msec/per pulse, 2 pulses with 1 sec interval; HeLa cells: 130V, 950 µF capacitance, exponential decay pulse.

### CRISPR-Cas9 mediated knockout and knockin

For UPF3B^KO#1^, two pX330 plasmids carrying two guide RNA sequences were co-transfected into HCT116 cells using JetOptimus (PolyPlus) as described above. After 2-3 weeks, single clones were isolated and screened for genomic deletion.

For UPF3A^KO#1^, UPF3B^KO#3^ and UPF3B^KO#4^, two guide RNAs per gene were synthesized (IDT or Synthego). 50 pmol of Cas9 recombinant protein (Berkeley Q3) and 60 pmol of each guide RNA were incubated for ∼20 mins in 10 µl reaction supplemented with Ingenio Electroporation Solution. 2.5 × 10^5^ cells were then mixed with CRISPR-Cas9 RNP complex and a final volume of 50 μl was used for the final electroporation reaction as described above. After 2-3 weeks, single clones were isolated and screened for genomic deletion.

For resistance marker-based knockouts, donor plasmids carrying antibiotic resistance genes and homology arms along with pX330 plasmid expressing guide RNAs that targeted Cas9 close to the insert site were co-transfected using JetOptimus (PolyPlus).

For knock-in of small affinity tags, HCT116 cells were synchronized using 2 μg/mL aphidicolin overnight and the synchronization was released 4 hours before the electroporation of CRISPR RNP complex. CRISPR RNP complex was prepared as described above and supplemented with 150 pmol of ssODN (50nt homology arms each side of the affinity tag). Electroporation was performed using Gene Pulser Xcell Electroporation System as described above.

For MYC-UPF2, we were unable to achieve efficient knock-in without selection. We used a resistance marker-based knock-in approach where a donor plasmid carrying hygromycin resistance marker-P2A-MYC tag in frame with UPF2 ORF was co-transfected with pX330 expressing guide RNA targeting a site close to the UPF2 start codon. Hygromycin resistant clones were isolated and screened for correct insert. All DNA sequences edited via CRISPR-Cas9 were confirmed by Sanger sequencing.

### RNA extraction

Cells were homogenized in TRI-reagent and RNA was extracted using one of three different methods. (1) RNA extraction was done following the manufacturer’s protocol, and then the extracted RNA was treated with 2 units of DNase I (NEB) and further cleaned up via phenol-chloroform (pH 4.3) extraction and standard ethanol precipitation. (2) One volume of ethanol was added to the TRI-reagent homogenized sample and the mixture was loaded onto a silica column (Epoch). The flowthrough was discarded and the column was washed once with high salt wash buffer (1.2 M Guanidine Thiocyanate; 10 mM Tris-HCl pH 7.0; 66% Ethanol) and twice with low salt wash buffer (10 mM Tris-HCl pH 7.0; 80% Ethanol). RNAs were eluted with water and subject to DNase I treatment as above. To the DNase I digested RNA, three volumes of RNA binding buffer (5.5 M Guanidine Thiocyanate; 0.55 M Sodium Acetate; 10 mM EDTA) and 4 volumes of ethanol were added. The mixture was then loaded into a silica column as above and washed twice with low salt wash buffer. RNA was then eluted in water. (3) All steps followed the second method with two exceptions: the silica columns were replaced with magnetic carboxylate modified beads (Cytiva) and the DNase I digestion was performed in the presence of beads (RNA gets eluted from beads upon addition of DNase I digestion mix). RNA was then re-bound to the magnetic beads by adding 5 volumes of ethanol. RNAs are then washed with low salt wash buffer twice before eluting in water.

### RT-qPCR

1.5 µg total RNA was reverse transcribed using Maxima RNaseH Minus Reverse Transcriptase following manufacturer’s protocol except that only 0.4 µl of the reverse transcriptase was added instead of 1 µl. cDNAs were then diluted to 5 ng/µl, and 3 µl of diluted cDNA was used per reaction. qPCR reactions were set up using iTaq Universal SYBR Green Supermix (Bio-Rad) with triplicates of 10 µl per reaction. qPCR was performed on a CFX-connect (Bio-Rad) equipment.

### β-globin reporter assays and Northern blots

For pulse-chase assays, 75,000 HeLa Tet-off cells were plated in each well of a 12-well plate. After 24h, reporter mRNA and protein expression plasmids were transfected using JetPrime, following manufacturer’s protocol (using 3:1 ratio of reagent to µg DNA). Cells were transfected with 200 ng pTet2 β39 plasmid, 20 ng β-GAP internal control, 10 ng pcDNA3ez-YFP, and 20 ng pcDNA3ez-FLAG-UPF3A/B. 100 ng/ml Tetracycline was used to suppress β39 expression. After 24h, tetracycline was removed to induce β39 expression overnight (∼16h). Tetracycline (1 µg/ml) was added, and cells were harvested in 0.5 ml TRIzol at indicated time points. For steady state assays, cells were transfected and induced in the same way as the reporter decay assay. At the day of harvesting, 1 µg/ml of tetracycline was added to all cells and cells are harvested 4-6 hours post transcription shut-off. RNA was extracted as above and Northern blotting was performed as described previously (Mabin et al., 2018).

### Protein Immunoprecipitation

Cells were washed with PBS and lysed with Gentle Hypotonic Lysis Buffer (20 mM Tris-HCl pH 7.5; 15 mM NaCl; 10 mM EDTA; 0.1% Triton X-100; 1× protease inhibitor cocktail; EDTA was replaced with 0.6 mM MgCl_2_ for magnesium-dependent IP (Fig. S6B)). A short 4-6 secs sonication pulse (10% amplitude) was applied to solubilize the chromatin fraction. 2-5 µl of the FLAG magnetic beads (Sigma) for FLAG-IP, and ∼1 µg primary antibody conjugated with protein-A dynabeads (Thermo Fisher) for EIF4A3-IP or CASC3-IP, were added to cell lysates and nutated at 4°C for 30-60 mins. Magnetic beads were then washed 8 times with Isotonic Wash Buffer (20 mM Tris-HCl pH 7.5; 150 mM NaCl; 0.1% IGEPAL CA-630). FLAG proteins were eluted for 10-20 mins with 250 ug/ml 3× FLAG peptide (APExBIO) in Isotonic Wash Buffer at 37°C. Primary antibody conjugated with protein-A beads were eluted for 5 mins in Clear Sample Buffer (100 mM Tris-HCl pH 6.8; 4% SDS; 10 mM EDTA) at 37°C.

### Total RNA-Seq library construction

800 ng of total RNA was rRNA-depleted using RiboCop rRNA Depletion Kit V1.3 (Lexogen) following manufacturer’s protocol. Libraries were then constructed using CORALL Total RNA-Seq Library Prep Kit (Lexogen). Libraries were quantified on RNA TapeStation and mixed at equimolar ratio for paired-end (2 × 150bp) sequencing using HiSeq4000 (Novogene) platform. Due to the relative short length of our libraries, we used only read 1 sequence for downstream analysis. We had three batches of experiments performed at different times, and only RNA-Seq samples sequenced at the same time were compared during the downstream analysis.

### Total RNA-Seq analysis

A reference script for mapping RNA-Seq libraries to the reference genome was kindly provided by Lexogen. For every fastq file, the first 10bp of each read (UMI) were extracted and appended to the header line using a custom Awk script and saved to a new file together with the remainder of the reads starting at position 13. Adapter trimming was then performed with cutadapt (Martin, 2011) for the sequence “AGATCGGAAGAGCACACGTCTGAACTCCAGTCAC”. Trimmed reads were aligned to the reference genome (GRCh38.p13) using STAR aligner (Dobin et al., 2013). Each output bam file was then indexed with samtools (Danecek et al., 2021) and deduplicated with UMI-tools (Smith et al., 2017) using the umi_tools dedup command with the [--method=unique -- multimapping-detection-method=NH] options. The fastq file with deduplicated reads was then extracted from the deduplicated bam file using samtools. Next, deduplicated fastq files served as input into pseudoalignment tool Kallisto (Bray et al., 2016) to quantify transcript abundance based on the Ensembl release 100 transcript reference. Tximport (Soneson et al., 2015) was used to extract transcript abundance from Kallisto results and generate count matrices for DESeq2. TPMs calculated from Kallisto results for each transcript were averaged for each experimental condition. We filtered out all the transcripts that have TPM less than 1 in all experimental conditions. After filtering out the transcripts, we use the RUVSeq (Risso et al., 2014) package [RUVs-method] to remove unwanted variation following the instruction manual. DESeq2 (Love et al., 2014) was then used to identify differentially expressed transcripts and calculate their fold changes.

### PTC+ and PTC-transcript lists

To generate a list of PTC+ transcripts and their PTC-counterparts, we used a custom Python script that takes all human transcripts in Ensembl annotation (version 100) along with their exon and 3′UTR coordinates to annotate each transcript as PTC+ if the 3′UTR begins more than 50 nt upstream of the exon junction or if there are more than one exon junctions downstream of the stop codon.

### RIPiT-Seq

RIPiT-Seq was performed as described (Yi & Singh, 2021). Four biological replicates each were performed for FLAG-MAGOH:EIF4A3, FLAG-UPF3B:EIF4A3, and FLAG-CASC3:EIF4A3 and sequenced on the HiSeq4000 (Novogene) platform.

### RIPiT-Seq quantification and differential occupancy analysis

Four replicates each of MAGOH-EJC, CASC3-EJC, and UPF3B-EJC RIPiT-Seq were obtained, for a total of 12 samples. RIPiT-Seq data analysis was performed similarly to our previous studies (Mabin et al., 2018; Patton et al., 2020). In short, the first 8 bp of each read (UMI) were extracted and appended to the header line using a custom Awk script and saved to a new file together with the remainder of the reads starting at position 9. Cutadapt (Martin, 2011) [--discard-untrimmed -g ^CC --no-indels * --discard-untrimmed -O 12 -a TGGAATTCTCGGGTGCCAAGG -] is used to retain any reads start with “CC” and ends with mirCat-33 adapter. Fastq files are further cleaned up by only retaining reads unable to align to a custom reference of abundant RNA sequences using STAR aligner [--outReadsUnmapped Fastx]. Trimmed reads were aligned to the reference genome (GRCh38.p13) using STAR aligner (Dobin et al., 2013). EJC signal for each gene was quantified using reads that overlap with the canonical EJC site (−39 to -9bp of 3′ end of non-last exon) and was averaged over all canonical EJC sites of a transcript (i.e., intron count). Any gene with EJC counts RPKM ≦ 5 was removed. Gene-level EJC signal was then input into DESeq2 for differential gene expression analysis (Love et al., 2014).

### Meta-exon analysis

RIPiT replicates and the exon annotation were used to compute total read depth as a function of distance from the 5’ start and 3’ end of each exon. Genes with less than 10 reads were discarded. Each remaining gene’s coverage distribution was normalized by the total number of reads of that gene and such normalized distributions were averaged across all genes. The average read distribution was then plotted with respect to the distance to the start or the end of the exon.

### Expression Normalized RIPiT Comparisons

Reads mapping to the canonical EJC region for each RIPiT-Seq sample were normalized by the total length of canonical region for each gene, and this length normalized EJC signal for each gene was divided by the RPKM of that gene from total RNA-seq. The resulting expression normalized signal for the CASC3, UPF3B, and MAGOH RIPiT-Seq were then correlated to the NMD efficiency of each gene that contains at least one PTC+ and one PTC-isoform. NMD efficiency of each gene is marked by the highest fold change of the PTC+ isoform in 3B^KO#1^:siUPF3A compared to WT:siNC.

## ACKNOWLEDGMENTS

We would like to thank Dr. Jens Lykke-Andersen for antibodies and plasmids, and Dr. Harold Fisk for kindly sharing the electroporation system with us. We would like to thank Dr. Niels Gehring for communicating unpublished results. Z.Y. is supported by Pelotonia Graduate Fellowship program and Center for RNA Biology Fellowship program at OSU. R.M.A. is supported by President’s Postdoctoral Scholars program and Pelotonia Postdoctoral Fellowship program at OSU. S.M. is supported by Department of Physics Undergraduate Summer Research Scholarship at OSU. C.N.D. is supported by Pelotonia Undergraduate Fellowship program and College of Arts and Sciences Undergraduate Research Scholarship at OSU. B.N.C. is supported by Center for RNA Biology Early-Stage Researcher Fellowship at OSU. This work is supported by NIH grant R01-GM120209 to G.S. We acknowledge an allocation of computation resources from the Ohio Supercomputer Center. Instrumentation was supported by NIH grant S10-OD023582.

## AUTHOR CONTRIBUTIONS

Conceptualization, Z.Y. and G.S.; Investigation, Z.Y., R.M.A., S.M., C.N.D., R.A.A., B.N.C., and R.D.P.; Writing - Original Draft, Z.Y. and G.S.; Writing - Review and Editing, Z.Y., R.M.A., S.M., R.D.P., R.B., and G.S.; Supervision, R.B. and G.S.

## DATA AVAILABILITY

RNA-Seq and RIPiT-Seq data are uploaded to GEO: GSE115977, which will be made accessible upon the acceptance of the manuscript.

## FIGURE LEGENDS

**Figure S1. Changes in gene expression and NMD upon *UPF3B* loss in HCT116 cells**.

A. Alteration in expression levels of known NMD genes in the two 3B^KO^ cell lines. RT-qPCR-based quantification of expression levels of previously characterized NMD-sensitive genes (x-axis) in the two 3B^KO^ HCT116 cell lines as compared to their levels in WT cells (set to 1). Relative levels from each replicate are shown by white circles. Error bars indicate standard error of means. The asterisk (*) represents p<0.05 in t-test with null hypothesis of true mean being 1 (n=3).

B. Immunoblot showing levels of UPF1 and UPF3B proteins in WT and 3B^KO#1^ cells (indicated on top) that were transfected with negative control (siNC) or UPF1-targeting (siUPF1) siRNAs. HNRNPA1 is a loading control.

C-E. MA plots showing differential transcript expression in RNA-Seq samples from (C) 3B^KO#1^ (siNC) versus WT (siNC), (D) UPF1-KD (siUPF1) versus control knockdown (siNC) in 3B^KO#1^, and (E) UPF1-KD (siUPF1) versus control knockdown (siNC) in WT cells. Each dot represents one transcript isoform with average read counts on the x axis and log_2_ fold change on the y axis. Transcripts that are significantly (adjusted p-value < 0.05) up (red)- or down (blue)-regulated >1.5-fold and their counts are indicated.

F. Isoform specific RT-qPCR measuring changes in levels of PTC+ and PTC-isoforms expressed from the indicated genes in WT and 3B^KO^ cells. Fold changes are with respect to the levels of PTC-isoforms in WT cells. Relative levels from each replicate are shown by white circles. Error bars indicate standard error of means. The asterisk (*) represents p<0.05 in t-test with null hypothesis of true mean being 1 (n=3).

G, H. CDF plots showing fold change in levels of transcripts with long, medium and short 3′ UTRs in (F) UPF1 (siUPF1) versus control (siNC) knockdown in WT cells, and (G) 3B^KO#1^ versus WT cells each transfected with control siRNA (siNC). p-values shown are from KS test comparing distribution of log_2_ fold changes in medium and long 3′UTR transcript groups as compared to the short 3′UTR transcript group. Number of transcripts in each group are also shown.

**Figure S2. UPF3A activates NMD in the absence of UPF3B**.

A. Western blots showing levels of EJC/UPF proteins or HNRNPA1 in input, normal rabbit IgG-IP or CASC3-IP fractions from WT and 3B^KO#1^ cells.

B. Western blots showing levels of EJC/UPF proteins or HNRNPA1 in input or FLAG-MAGOH followed by MYC-UPF2 tandem-IP fractions from WT, 3A^KO#2^, and 3B^KO#2^ cells. Samples were RNase A treated during the FLAG IP. The asterisk (*) represents the mouse heavy chain of the MYC-tag antibody.

C. Immunoblot showing levels of UPF3A and UPF3B proteins in WT and 3B^KO#1^ cells (indicated on top) that were transfected with negative control (siNC) or UPF3A-targeting (siUPF3A) siRNAs. HNRNPA1 is a loading control.

D. MA plots showing differential transcript expression in RNA-Seq samples from UPF3A-KD (siUPF3A) versus control knockdown (siNC) in WT HCT116 cells. Each dot represents one transcript isoform with average read counts on the x-axis and log_2_ fold change on the y-axis. Red and blue dots represent >1.5-fold up- or down-regulated transcripts, respectively, that are significantly changed (adjusted p-value < 0.05).

E. Schematic of *UPF3A* knockout (UPF3B-KO) strategies using CRISPR-Cas9. *UPF3A* locus is in black where rectangles represent exons and horizontal line denotes introns; coding region is shown as wider rectangles. Red arrowheads represent guide RNA targeting sites. In 3A^KO#1^ (top), two guide RNAs delete first and the second exons of UPF3A protein coding region. In 3A^KO#2^ (bottom), a donor template is used to insert blasticidin resistant gene (BlasticidinR) and Simian Virus 40 (SV40) polyadenylation signal at the cut site.

F. Immunoblot of UPF3A and UPF3B proteins in WT, 3A^KO^, 3B^KO^, and 3^DKO^ cells. EIF4A3 is used as a loading control.

G. Isoform specific RT-qPCR of PTC+ and PTC-isoforms from the indicated genes in WT, 3A^KO^, 3B^KO^, and 3^DKO^ cells. Relative levels from each replicate are shown by white circles Error bars indicate standard error of means. The asterisk (*) represents p<0.05 in t-test with null hypothesis of true mean being 1 (n=3).

H, I. CDF plots showing fold change in levels of transcripts with short, medium and long 3′UTRs in UPF3A (siUPF3A) versus control (siNC) knockdown in (F) WT cells, and (G) 3B^KO#1^ cells.

**Figure S3. CASC3 regulates UPF3-dependent NMD**.

A. Schematic of *UPF3B* knockout in HeLa Tet-off cells using CRISPR-Cas9. Red arrows represent two guide RNA targeting sites which will lead to the deletion of the first exon.

B. Protein immunoblot of UPF3B protein in WT and 3B^KO^ HeLa cells. HNRNPA1 is a loading control.

C. Western blots showing proteins on the right in input and EIF4A3-IP from WT and 3B^KO^ HeLa cells. Normal rabbit IgG is used for control IP.

D. Western blots as in C from HeLa WT and 3B^KO^ cells transfected with either siNC or siCASC3.

E. Meta-exon plot of MAGOH:EIF4A3, UPF3B:EIF4A3, and CASC3:EIF4A3 RIPiT-Seq read-distribution in the 100 nt region from the exon 5′ end.

F, G. Venn diagram of significantly enriched/depleted genes in CASC3:EIF4A3 or UPF3B:EIF4A3 RIPiT-Seq samples as compared to MAGOH:EIF4A3 RIPiT-Seq.

H, I. Scatter plots showing a comparison between gene-level EJC occupancy and NMD efficiency. UPF3B:EIF3A3 (H), or MAGOH:EIF4A3 (I) RIPiT-Seq signal normalized to expression level of individual genes is on the x-axis and NMD efficiency of each gene on the y-axis. For each gene in this analysis, NMD efficiency is the highest fold change (in 3B^KO#1^ siUPF3A to WT siNC) observed for its PTC+ isoform. R^2^ from the linear regression fit is shown on the top left.

J. Protein immunoblot of CASC3 protein in WT and CASC-KO HeLa cells. HNRNPA1 is a loading control.

**Figure S4. NMD in the absence of both UPF3 paralogs**.

A. Immunoblots showing levels of UPF1 and UPF2 proteins in 3^DKO#2^ cells that were transfected with negative control (siNC), UPF1-targeting (siUPF1), or UPF2-targeting (siUPF2) siRNAs. HNRNPA1 is a loading control.

B, C. MA plots showing transcript-level changes upon UPF2 (siUPF2) knockdown as compared to control knockdown (siNC) in, (B) WT cells, and (C) 3^DKO#2^ cells. Each dot represents one transcript with average read counts on the x-axis and log_2_ fold change on the y-axis. Red and blue dots represent transcripts up- or down-regulated more than 1.5-fold with an adjusted p-value < 0.05; these counts are shown on each plot.

**Figure S5. UPF3 paralogs differ in NMD activity**.

A, B. Protein immunoblots of FLAG-IP from 3^DKO#2^ cells expressing different FLAG-tagged human UPF3 proteins or their chimeras using Tet-on 3G system. HNRNPA1 is used as loading control and RNase A digestion control. FLAG-EGFP is used as an IP control.

**Figure S6. EJC interaction ability of human and mouse UPF3A proteins**.

Protein immunoblots of input and FLAG-IP from WT and 3^DKO#2^ cells FLAG-tagged human or mouse UPF3A as indicated above the lanes. HNRNPA1 is a loading and RNase A digestion control.

## REFERENCES

Addington, A. M., Gauthier, J., Piton, A., Hamdan, F. F., Raymond, A., Gogtay, N., Miller, R., Tossell, J., Bakalar, J., Germain, G., Gochman, P., Long, R., Rapoport, J. L., & Rouleau, G. A. (2011). A novel frameshift mutation in UPF3B identified in brothers affected with childhood onset schizophrenia and autism spectrum disorders. Molecular Psychiatry, 16(3), 238–239. https://doi.org/10.1038/mp.2010.59

Alrahbeni, T., Sartor, F., Anderson, J., Miedzybrodzka, Z., McCaig, C., & Müller, B. (2015). Full UPF3B function is critical for neuronal differentiation of neural stem cells. Molecular Brain, 8(1), 33. https://doi.org/10.1186/s13041-015-0122-1

Amrani, N., Ganesan, R., Kervestin, S., Mangus, D. A., Ghosh, S., & Jacobson, A. (2004). A faux 3’-UTR promotes aberrant termination and triggers nonsense-mediated mRNA decay. Nature, 432(7013), 112–118. https://doi.org/10.1038/nature03060

Avery, P., Vicente-Crespo, M., Francis, D., Nashchekina, O., Alonso, C. R., & Palacios, I. M. (2011). Drosophila Upf1 and Upf2 loss of function inhibits cell growth and causes animal death in a Upf3-independent manner. RNA, 17(4), 624–638. https://doi.org/10.1261/rna.2404211

Baird, T. D., Cheng, K. C.-C., Chen, Y.-C., Buehler, E., Martin, S. E., Inglese, J., & Hogg, J. R. (2018). ICE1 promotes the link between splicing and nonsense-mediated mRNA decay. ELife, 7, e33178. https://doi.org/10.7554/eLife.33178

Ballut, L., Marchadier, B., Baguet, A., Tomasetto, C., Séraphin, B., & Le Hir, H. (2005). The exon junction core complex is locked onto RNA by inhibition of eIF4AIII ATPase activity. Nature Structural & Molecular Biology, 12(10), 861–869. https://doi.org/10.1038/nsmb990

Behm-Ansmant, I., Gatfield, D., Rehwinkel, J., Hilgers, V., & Izaurralde, E. (2007). A conserved role for cytoplasmic poly(A)-binding protein 1 (PABPC1) in nonsense-mediated mRNA decay. The EMBO Journal, 26(6), 1591–1601. https://doi.org/10.1038/sj.emboj.7601588

Boehm, V., & Gehring, N. H. (2016). Exon Junction Complexes: Supervising the Gene Expression Assembly Line. Trends in Genetics, 32(11), 724–735. https://doi.org/10.1016/j.tig.2016.09.003

Bray, N. L., Pimentel, H., Melsted, P., & Pachter, L. (2016). Near-optimal probabilistic RNA-seq quantification. Nature Biotechnology, 34(5), 525–527. https://doi.org/10.1038/nbt.3519

Bruno, I. G., Karam, R., Huang, L., Bhardwaj, A., Lou, C. H., Shum, E. Y., Song, H.-W., Corbett, M. A., Gifford, W. D., Gecz, J., Pfaff, S. L., & Wilkinson, M. F. (2011). Identification of a MicroRNA that Activates Gene Expression by Repressing Nonsense-Mediated RNA Decay. Molecular Cell, 42(4), 500–510. https://doi.org/10.1016/j.molcel.2011.04.018

Buchwald, G., Ebert, J., Basquin, C., Sauliere, J., Jayachandran, U., Bono, F., Hir, H. L., & Conti, E. (2010). Insights into the recruitment of the NMD machinery from the crystal structure of a core EJC-UPF3b complex. Proceedings of the National Academy of Sciences, 107(22), 10050–10055. https://doi.org/10.1073/pnas.1000993107

Celik, A., Baker, R., He, F., & Jacobson, A. (2017). High-resolution profiling of NMD targets in yeast reveals translational fidelity as a basis for substrate selection. RNA, 23(5), 735–748. https://doi.org/10.1261/rna.060541.116

Chamieh, H., Ballut, L., Bonneau, F., & Le Hir, H. (2008). NMD factors UPF2 and UPF3 bridge UPF1 to the exon junction complex and stimulate its RNA helicase activity. Nature Structural & Molecular Biology, 15(1), 85–93. https://doi.org/10.1038/nsmb1330

Chan, W.-K., Bhalla, A. D., Le Hir, H., Nguyen, L. S., Huang, L., Gécz, J., & Wilkinson, M. F. (2009). A UPF3-mediated regulatory switch that maintains RNA surveillance. Nature Structural & Molecular Biology, 16(7), 747–753. https://doi.org/10.1038/nsmb.1612

Chan, W.-K., Huang, L., Gudikote, J. P., Chang, Y.-F., Imam, J. S., MacLean II, J. A., & Wilkinson, M. F. (2007). An alternative branch of the nonsense-mediated decay pathway. The EMBO Journal, 26(7), 1820–1830. https://doi.org/10.1038/sj.emboj.7601628

Danecek, P., Bonfield, J. K., Liddle, J., Marshall, J., Ohan, V., Pollard, M. O., Whitwham, A., Keane, T., McCarthy, S. A., Davies, R. M., & Li, H. (2021). Twelve years of SAMtools and BCFtools. GigaScience, 10(2). https://doi.org/10.1093/gigascience/giab008

Dobin, A., Davis, C. A., Schlesinger, F., Drenkow, J., Zaleski, C., Jha, S., Batut, P., Chaisson, M., & Gingeras, T. R. (2013). STAR: Ultrafast universal RNA-seq aligner. Bioinformatics, 29(1), 15–21. https://doi.org/10.1093/bioinformatics/bts635

Eberle, A. B., Stalder, L., Mathys, H., Orozco, R. Z., & Mühlemann, O. (2008). Posttranscriptional Gene Regulation by Spatial Rearrangement of the 3’ Untranslated Region. PLOS Biology, 6(4), e92. https://doi.org/10.1371/journal.pbio.0060092

Gehring, N. H., Kunz, J. B., Neu-Yilik, G., Breit, S., Viegas, M. H., Hentze, M. W., & Kulozik, A. E. (2005). Exon-Junction Complex Components Specify Distinct Routes of Nonsense-Mediated mRNA Decay with Differential Cofactor Requirements. Molecular Cell, 20(1), 65–75. https://doi.org/10.1016/j.molcel.2005.08.012

Gerbracht, J. V., Boehm, V., Britto-Borges, T., Kallabis, S., Wiederstein, J. L., Ciriello, S., Aschemeier, D. U., Krüger, M., Frese, C. K., Altmüller, J., Dieterich, C., & Gehring, N. H. (2020). CASC3 promotes transcriptome-wide activation of nonsense-mediated decay by the exon junction complex. Nucleic Acids Research, 48(15), 8626–8644. https://doi.org/10.1093/nar/gkaa564

He, F., Brown, A. H., & Jacobson, A. (1997). Upf1p, Nmd2p, and Upf3p are interacting components of the yeast nonsense-mediated mRNA decay pathway. Molecular and Cellular Biology, 17(3), 1580–1594. https://doi.org/10.1128/MCB.17.3.1580

He, F., & Jacobson, A. (2015). Nonsense-Mediated mRNA Decay: Degradation of Defective Transcripts Is Only Part of the Story. Annual Review of Genetics, 49(1), 339–366. https://doi.org/10.1146/annurev-genet-112414-054639

Hir, H. L., Saulière, J., & Wang, Z. (2016). The exon junction complex as a node of post-transcriptional networks. Nature Reviews Molecular Cell Biology, 17(1), 41–54. https://doi.org/10.1038/nrm.2015.7

Hogg, J. R., & Goff, S. P. (2010). Upf1 Senses 3’UTR Length to Potentiate mRNA Decay. Cell, 143(3), 379–389. https://doi.org/10.1016/j.cell.2010.10.005

Huang, L., Lou, C.-H., Chan, W., Shum, E. Y., Shao, A., Stone, E., Karam, R., Song, H.-W., & Wilkinson, M. F. (2011). RNA Homeostasis Governed by Cell Type-Specific and Branched Feedback Loops Acting on NMD. Molecular Cell, 43(6), 950–961. https://doi.org/10.1016/j.molcel.2011.06.031

Huang, L., Shum, E. Y., Jones, S. H., Lou, C.-H., Dumdie, J., Kim, H., Roberts, A. J., Jolly, L. A., Espinoza, J. L., Skarbrevik, D. M., Phan, M. H., Cook-Andersen, H., Swerdlow, N. R., Gecz, J., & Wilkinson, M. F. (2018). A Upf3b-mutant mouse model with behavioral and neurogenesis defects. Molecular Psychiatry, 23(8), 1773–1786. https://doi.org/10.1038/mp.2017.173

Ivanov, P. V., Gehring, N. H., Kunz, J. B., Hentze, M. W., & Kulozik, A. E. (2008). Interactions between UPF1, eRFs, PABP and the exon junction complex suggest an integrated model for mammalian NMD pathways. The EMBO Journal, 27(5), 736–747. https://doi.org/10.1038/emboj.2008.17

Kadlec, J., Izaurralde, E., & Cusack, S. (2004). The structural basis for the interaction between nonsense-mediated mRNA decay factors UPF2 and UPF3. Nature Structural & Molecular Biology, 11(4), 330–337. https://doi.org/10.1038/nsmb741

Karousis, E. D., Gurzeler, L.-A., Annibaldis, G., Dreos, R., & Mühlemann, O. (2020). Human NMD ensues independently of stable ribosome stalling. Nature Communications, 11(1), 4134. https://doi.org/10.1038/s41467-020-17974-z

Karousis, E. D., & Mühlemann, O. (2019). Nonsense-Mediated mRNA Decay Begins Where Translation Ends. Cold Spring Harbor Perspectives in Biology, 11(2), a032862. https://doi.org/10.1101/cshperspect.a032862

Kashima, I., Yamashita, A., Izumi, N., Kataoka, N., Morishita, R., Hoshino, S., Ohno, M., Dreyfuss, G., & Ohno, S. (2006). Binding of a novel SMG-1–Upf1–eRF1–eRF3 complex (SURF) to the exon junction complex triggers Upf1 phosphorylation and nonsense-mediated mRNA decay. Genes & Development, 20(3), 355–367. https://doi.org/10.1101/gad.1389006

Kishor, A., Fritz, S. E., & Hogg, J. R. (2019). Nonsense-mediated mRNA decay: The challenge of telling right from wrong in a complex transcriptome. WIREs RNA, 10(6), e1548. https://doi.org/10.1002/wrna.1548

Kunz, J. B., Neu-Yilik, G., Hentze, M. W., Kulozik, A. E., & Gehring, N. H. (2006). Functions of hUpf3a and hUpf3b in nonsense-mediated mRNA decay and translation. RNA, 12(6), 1015–1022. https://doi.org/10.1261/rna.12506

Kurosaki, T., Popp, M. W., & Maquat, L. E. (2019). Quality and quantity control of gene expression by nonsense-mediated mRNA decay. Nature Reviews Molecular Cell Biology, 20(7), 406–420. https://doi.org/10.1038/s41580-019-0126-2

Laumonnier, F., Shoubridge, C., Antar, C., Nguyen, L. S., Van Esch, H., Kleefstra, T., Briault, S., Fryns, J. P., Hamel, B., Chelly, J., Ropers, H. H., Ronce, N., Blesson, S., Moraine, C., Gécz, J., & Raynaud, M. (2010). Mutations of the UPF3B gene, which encodes a protein widely expressed in neurons, are associated with nonspecific mental retardation with or without autism. Molecular Psychiatry, 15(7), 767–776. https://doi.org/10.1038/mp.2009.14

López-Perrote, A., Castaño, R., Melero, R., Zamarro, T., Kurosawa, H., Ohnishi, T., Uchiyama, A., Aoyagi, K., Buchwald, G., Kataoka, N., Yamashita, A., & Llorca, O. (2016). Human nonsense-mediated mRNA decay factor UPF2 interacts directly with eRF3 and the SURF complex. Nucleic Acids Research, 44(4), 1909–1923. https://doi.org/10.1093/nar/gkv1527

Love, M. I., Huber, W., & Anders, S. (2014). Moderated estimation of fold change and dispersion for RNA-seq data with DESeq2. Genome Biology, 15(12), 550. https://doi.org/10.1186/s13059-014-0550-8

Lykke-Andersen, J., Shu, M.-D., & Steitz, J. A. (2000). Human Upf Proteins Target an mRNA for Nonsense-Mediated Decay When Bound Downstream of a Termination Codon. Cell, 103(7), 1121–1131. https://doi.org/10.1016/S0092-8674(00)00214-2

Lynch, S. A., Nguyen, L. S., Ng, L. Y., Waldron, M., McDonald, D., & Gecz, J. (2012). Broadening the phenotype associated with mutations in UPF3B: Two further cases with renal dysplasia and variable developmental delay. European Journal of Medical Genetics, 55(8), 476–479. https://doi.org/10.1016/j.ejmg.2012.03.010

Ma, Z., Zhu, P., Shi, H., Guo, L., Zhang, Q., Chen, Y., Chen, S., Zhang, Z., Peng, J., & Chen, J. (2019). PTC-bearing mRNA elicits a genetic compensation response via Upf3a and COMPASS components. Nature, 568(7751), 259–263. https://doi.org/10.1038/s41586-019-1057-y

Mabin, J. W., Woodward, L. A., Patton, R. D., Yi, Z., Jia, M., Wysocki, V. H., Bundschuh, R., & Singh, G. (2018). The Exon Junction Complex Undergoes a Compositional Switch that Alters mRNP Structure and Nonsense-Mediated mRNA Decay Activity. Cell Reports, 25(9), 2431–2446.e7. https://doi.org/10.1016/j.celrep.2018.11.046

Martin, M. (2011). Cutadapt removes adapter sequences from high-throughput sequencing reads. EMBnet. Journal, 17(1), 10–12. https://doi.org/10.14806/ej.17.1.200

Medghalchi, S. M., Frischmeyer, P. A., Mendell, J. T., Kelly, A. G., Lawler, A. M., & Dietz, H. C. (2001). Rent1, a trans-effector of nonsense-mediated mRNA decay, is essential for mammalian embryonic viability. Human Molecular Genetics, 10(2), 99–105. https://doi.org/10.1093/hmg/10.2.99

Melero, R., Buchwald, G., Castaño, R., Raabe, M., Gil, D., Lázaro, M., Urlaub, H., Conti, E., & Llorca, O. (2012). The cryo-EM structure of the UPF–EJC complex shows UPF1 poised toward the RNA 3’ end. Nature Structural & Molecular Biology, 19(5), 498–505. https://doi.org/10.1038/nsmb.2287

Mendell, J. T., Sharifi, N. A., Meyers, J. L., Martinez-Murillo, F., & Dietz, H. C. (2004). Nonsense surveillance regulates expression of diverse classes of mammalian transcripts and mutes genomic noise. Nature Genetics, 36(10), 1073–1078. https://doi.org/10.1038/ng1429

Natsume, T., Kiyomitsu, T., Saga, Y., & Kanemaki, M. T. (2016). Rapid Protein Depletion in Human Cells by Auxin-Inducible Degron Tagging with Short Homology Donors. Cell Reports, 15(1), 210–218. https://doi.org/10.1016/j.celrep.2016.03.001

Neu-Yilik, G., Raimondeau, E., Eliseev, B., Yeramala, L., Amthor, B., Deniaud, A., Huard, K., Kerschgens, K., Hentze, M. W., Schaffitzel, C., & Kulozik, A. E. (2017). Dual function of UPF3B in early and late translation termination. The EMBO Journal, 36(20), 2968–2986. https://doi.org/10.15252/embj.201797079

Nguyen, L. S., Jolly, L., Shoubridge, C., Chan, W. K., Huang, L., Laumonnier, F., Raynaud, M., Hackett, A., Field, M., Rodriguez, J., Srivastava, A. K., Lee, Y., Long, R., Addington, A. M., Rapoport, J. L., Suren, S., Hahn, C. N., Gamble, J., Wilkinson, M. F., … Gecz, J. (2012). Transcriptome profiling of UPF3B/NMD-deficient lymphoblastoid cells from patients with various forms of intellectual disability. Molecular Psychiatry, 17(11), 1103–1115. https://doi.org/10.1038/mp.2011.163

Patton, R. D., Sanjeev, M., Woodward, L. A., Mabin, J. W., Bundschuh, R., & Singh, G. (2020). Chemical crosslinking enhances RNA immunoprecipitation for efficient identification of binding sites of proteins that photo-crosslink poorly with RNA. RNA, rna.074856.120. https://doi.org/10.1261/rna.074856.120

Peixeiro, I., Inácio, Â., Barbosa, C., Silva, A. L., Liebhaber, S. A., & Romão, L. (2012). Interaction of PABPC1 with the translation initiation complex is critical to the NMD resistance of AUG-proximal nonsense mutations. Nucleic Acids Research, 40(3), 1160–1173. https://doi.org/10.1093/nar/gkr820

Ran, F. A., Hsu, P. D., Wright, J., Agarwala, V., Scott, D. A., & Zhang, F. (2013). Genome engineering using the CRISPR-Cas9 system. Nature Protocols, 8(11), 2281–2308. https://doi.org/10.1038/nprot.2013.143

Risso, D., Ngai, J., Speed, T. P., & Dudoit, S. (2014). Normalization of RNA-seq data using factor analysis of control genes or samples. Nature Biotechnology, 32(9), 896–902. https://doi.org/10.1038/nbt.2931

Shum, E. Y., Jones, S. H., Shao, A., Dumdie, J., Krause, M. D., Chan, W.-K., Lou, C.-H., Espinoza, J. L., Song, H.-W., Phan, M. H., Ramaiah, M., Huang, L., McCarrey, J. R., Peterson, K. J., De Rooij, D. G., Cook-Andersen, H., & Wilkinson, M. F. (2016). The Antagonistic Gene Paralogs Upf3a and Upf3b Govern Nonsense-Mediated RNA Decay. Cell, 165(2), 382–395. https://doi.org/10.1016/j.cell.2016.02.046

Singh, G., Jakob, S., Kleedehn, M. G., & Lykke-Andersen, J. (2007). Communication with the Exon-Junction Complex and Activation of Nonsense-Mediated Decay by Human Upf Proteins Occur in the Cytoplasm. Molecular Cell, 27(5), 780–792. https://doi.org/10.1016/j.molcel.2007.06.030

Singh, G., Kucukural, A., Cenik, C., Leszyk, J. D., Shaffer, S. A., Weng, Z., & Moore, M. J. (2012). The Cellular EJC Interactome Reveals Higher-Order mRNP Structure and an EJC-SR Protein Nexus. Cell, 151(4), 750–764. https://doi.org/10.1016/j.cell.2012.10.007

Singh, G., Rebbapragada, I., & Lykke-Andersen, J. (2008). A Competition between Stimulators and Antagonists of Upf Complex Recruitment Governs Human Nonsense-Mediated mRNA Decay. PLOS Biology, 6(4), e111. https://doi.org/10.1371/journal.pbio.0060111

Smith, T., Heger, A., & Sudbery, I. (2017). UMI-tools: Modeling sequencing errors in Unique Molecular Identifiers to improve quantification accuracy. Genome Research, 27(3), 491–499. https://doi.org/10.1101/gr.209601.116

Soneson, C., Love, M. I., & Robinson, M. D. (2015). Differential analyses for RNA-seq: Transcript-level estimates improve gene-level inferences. F1000Research, 4, 1521. https://doi.org/10.12688/f1000research.7563.1

Tani, H., Imamachi, N., Salam, K. A., Mizutani, R., Ijiri, K., Irie, T., Yada, T., Suzuki, Y., & Akimitsu, N. (2012). Identification of hundreds of novel UPF1 target transcripts by direct determination of whole transcriptome stability. RNA Biology, 9(11), 1370–1379. https://doi.org/10.4161/rna.22360

Tarpey, P. S., Lucy Raymond, F., Nguyen, L. S., Rodriguez, J., Hackett, A., Vandeleur, L., Smith, R., Shoubridge, C., Edkins, S., Stevens, C., O’Meara, S., Tofts, C., Barthorpe, S., Buck, G., Cole, J., Halliday, K., Hills, K., Jones, D., Mironenko, T., … Gécz, J. (2007). Mutations in UPF3B, a member of the nonsense-mediated mRNA decay complex, cause syndromic and nonsyndromic mental retardation. Nature Genetics, 39(9), 1127–1133. https://doi.org/10.1038/ng2100

Turner, J. M. A. (2007). Meiotic sex chromosome inactivation. Development, 134(10), 1823–1831. https://doi.org/10.1242/dev.000018

Wallmeroth, D., Boehm, V., Lackmann, J.-W., Altmueller, J., Dieterich, C., & Gehring, N. H. (2021). UPF3A and UPF3B are redundant and modular activators of nonsense-mediated mRNA decay in human cells. BioRxiv, 2021.07.07.451444. https://doi.org/10.1101/2021.07.07.451444

Wang, W., Czaplinski, K., Rao, Y., & Peltz, S. W. (2001). The role of Upf proteins in modulating the translation read-through of nonsense-containing transcripts. The EMBO Journal, 20(4), 880–890. https://doi.org/10.1093/emboj/20.4.880

Weischenfeldt, J., Damgaard, I., Bryder, D., Theilgaard-Mönch, K., Thoren, L. A., Nielsen, F. C., Jacobsen, S. E. W., Nerlov, C., & Porse, B. T. (2008). NMD is essential for hematopoietic stem and progenitor cells and for eliminating by-products of programmed DNA rearrangements. Genes & Development, 22(10), 1381–1396. https://doi.org/10.1101/gad.468808

Wittmann, J., Hol, E. M., & Jäck, H.-M. (2006). HUPF2 Silencing Identifies Physiologic Substrates of Mammalian Nonsense-Mediated mRNA Decay. Molecular and Cellular Biology, 26(4), 1272–1287. https://doi.org/10.1128/MCB.26.4.1272-1287.2006

Woodward, L. A., Mabin, J. W., Gangras, P., & Singh, G. (2017). The exon junction complex: A lifelong guardian of mRNA fate. WIREs RNA, 8(3), e1411. https://doi.org/10.1002/wrna.1411

Xu, X., Zhang, L., Tong, P., Xun, G., Su, W., Xiong, Z., Zhu, T., Zheng, Y., Luo, S., Pan, Y., Xia, K., & Hu, Z. (2013). Exome sequencing identifies UPF3B as the causative gene for a Chinese non-syndrome mental retardation pedigree. Clinical Genetics, 83(6), 560–564. https://doi.org/10.1111/cge.12014

Yi, Z., Sanjeev, M., & Singh, G. (2021). The Branched Nature of the Nonsense-Mediated mRNA Decay Pathway. Trends in Genetics, 37(2), 143–159. https://doi.org/10.1016/j.tig.2020.08.010

Yi, Z., & Singh, G. (2021). Chapter Seventeen - RIPiT-Seq: A tandem immunoprecipitation approach to reveal global binding landscape of multisubunit ribonucleoproteins. In B. Tian (Ed.), Methods in Enzymology (Vol. 655, pp. 401–425). Academic Press. https://doi.org/10.1016/bs.mie.2021.03.019

Yusa, K., Zhou, L., Li, M. A., Bradley, A., & Craig, N. L. (2011). A hyperactive piggyBac transposase for mammalian applications. Proceedings of the National Academy of Sciences, 108(4), 1531–1536. https://doi.org/10.1073/pnas.1008322108

