## Supplement Figures S1-S6 for "Mammalian UPF3A and UPF3B activate NMD independently of their EJC binding"

### Figure S1

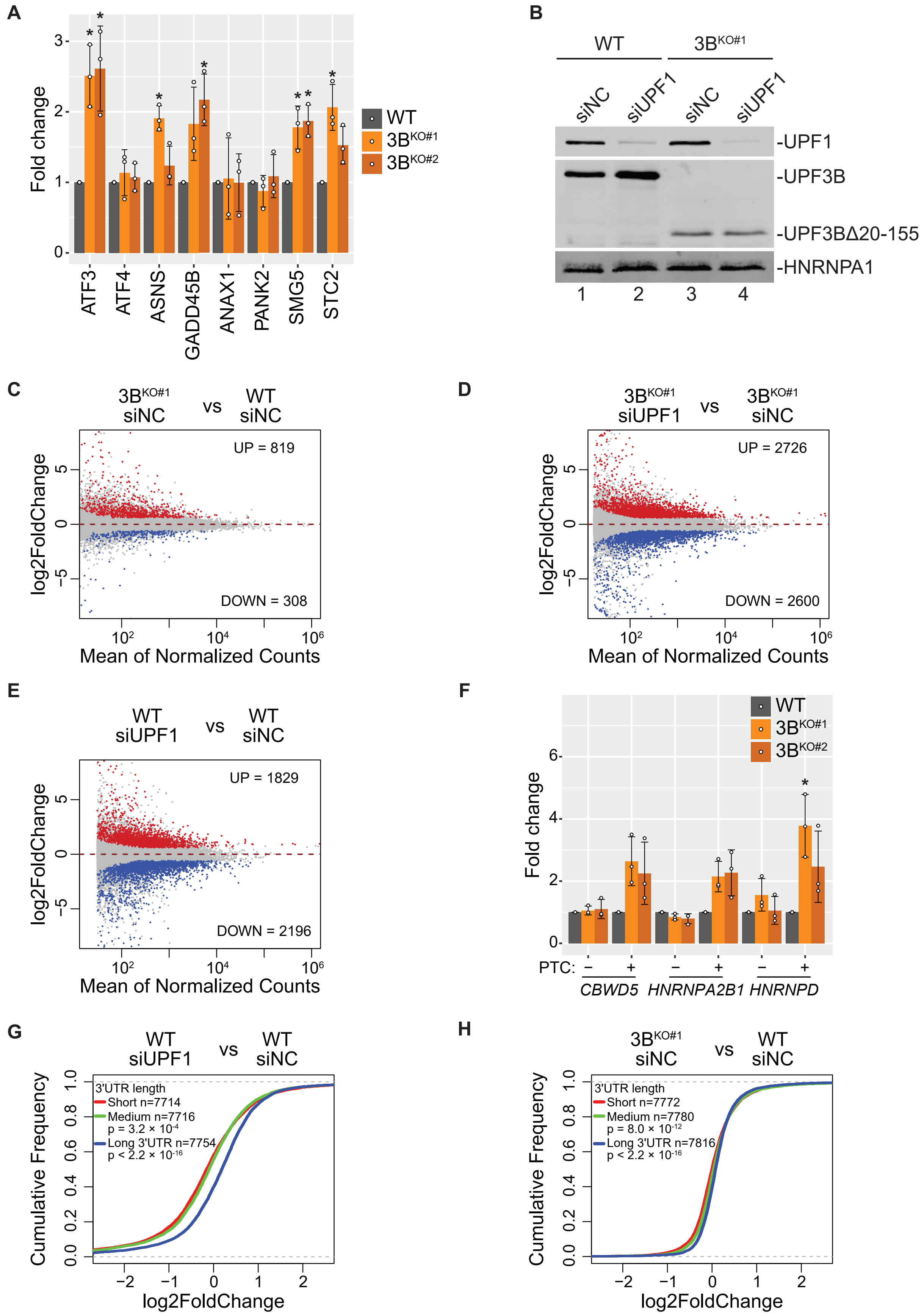


**Figure S1. Changes in gene expression and NMD upon UPF3B loss in HCT116 cells.**
A. Alteration in expression levels of known NMD genes in the two $3B^{KO}$ cell lines. RT-qPCR-based quantification of expression levels of previously characterized NMD-sensitive genes (x-axis) in the two $3B^{KO}$ HCT116 cell lines as compared to their levels in WT cells (set to 1). Relative levels from each replicate are shown by white circles. Error bars indicate standard error of means. The asterisk (*) represents p<0.05 in t-test with null hypothesis of true mean being 1 (n=3).
B. Immunoblot showing levels of UPF1 and UPF3B proteins in WT and $3B^{KO\#1}$ cells (indicated on top) that were transfected with negative control (siNC) or UPF1-targeting (siUPF1) siRNAs. HNRNPA1 is a loading control.
C-E. MA plots showing differential transcript expression in RNA-Seq samples from (C) $3B^{KO\#1}$ (siNC) versus WT (siNC), (D) UPF1-KD (siUPF1) versus control knockdown (siNC) in $3B^{KO\#1}$, and (E) UPF1-KD (siUPF1) versus control knockdown (siNC) in WT cells. Each dot represents one transcript isoform with average read counts on the x axis and log2 fold change on the y axis. Transcripts that are significantly (adjusted p-value < 0.05) up (red)- or down (blue)- regulated >1.5-fold and their counts are indicated.
F. Isoform specific RT-qPCR measuring changes in levels of PTC+ and PTC- isoforms expressed from the indicated genes in WT and $3B^{KO}$ cells. Fold changes are with respect to the levels of PTC- isoforms in WT cells. Relative levels from each replicate are shown by white circles. Error bars indicate standard error of means. The asterisk (*) represents p<0.05 in t-test with null hypothesis of true mean being 1 (n=3).
G, H. CDF plots showing fold change in levels of transcripts with long, medium and short 3′ UTRs in (F) UPF1 (siUPF1) versus control (siNC) knockdown in WT cells, and (G) $3B^{KO\#1}$ versus WT cells each transfected with control siRNA (siNC). p-values shown are from KS test comparing distribution of log2 fold changes in medium and long 3′UTR transcript groups as compared to the short 3′UTR transcript group. Number of transcripts in each group are also shown.

### Figure S2

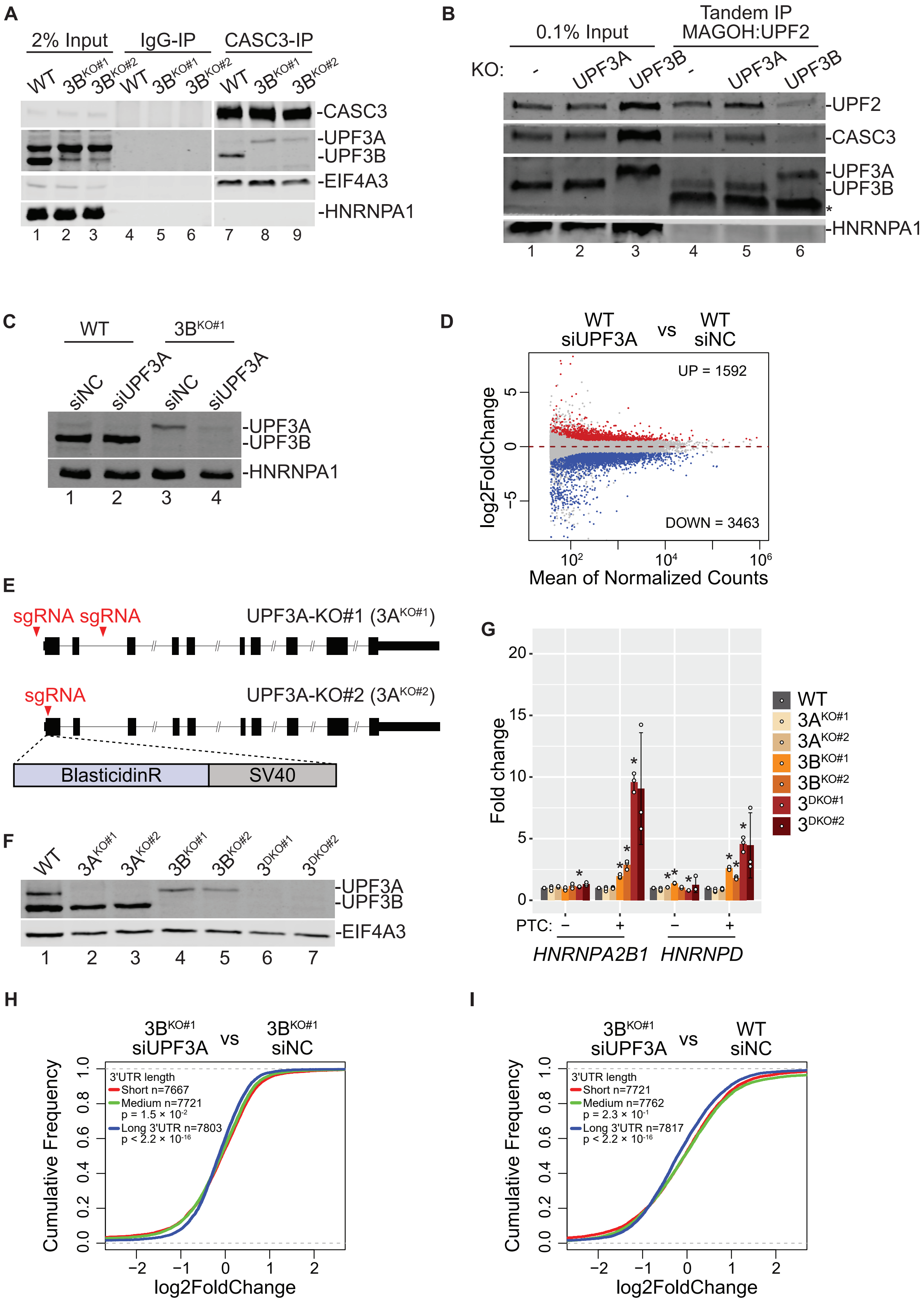


**Figure S2. UPF3A activates NMD in the absence of UPF3B.**
A. Western blots showing levels of EJC/UPF proteins or HNRNPA1 in input, normal rabbit IgG-IP or CASC3-IP fractions from WT and $3B^{KO\#1}$ cells.
B. Western blots showing levels of EJC/UPF proteins or HNRNPA1 in input or FLAG-MAGOH followed by MYC-UPF2 tandem-IP fractions from WT, $3A^{KO\#2}$, and $3B^{KO\#2}$ cells. Samples were RNase A treated during the FLAG IP. The asterisk (*) represents the mouse heavy chain of the MYC-tag antibody.
C. Immunoblot showing levels of UPF3A and UPF3B proteins in WT and $3B^{KO\#1}$ cells (indicated on top) that were transfected with negative control (siNC) or UPF3A-targeting (siUPF3A) siRNAs. HNRNPA1 is a loading control.
D. MA plots showing differential transcript expression in RNA-Seq samples from UPF3A-KD (siUPF3A) versus control knockdown (siNC) in WT HCT116 cells. Each dot represents one transcript isoform with average read counts on the x-axis and log2 fold change on the y-axis. Red and blue dots represent >1.5-fold up- or down- regulated transcripts, respectively, that are significantly changed (adjusted p-value < 0.05).
E. Schematic of UPF3A knockout (UPF3A-KO) strategies using CRISPR-Cas9. UPF3A locus is in black where rectangles represent exons and horizontal line denotes introns; coding region is shown as wider rectangles. Red arrowheads represent guide RNA targeting sites. In $3A^{KO\#1}$ (top), two guide RNAs delete first and the second exons of UPF3A protein coding region. In $3A^{KO\#2}$ (bottom), a donor template is used to insert blasticidin resistant gene (BlasticidinR) and Simian Virus 40 (SV40) polyadenylation signal at the cut site.
F. Immunoblot of UPF3A and UPF3B proteins in WT, $3A^{KO}$, $3B^{KO}$, and $3^{DKO}$ cells. EIF4A3 is used as a loading control.

### Figure S3

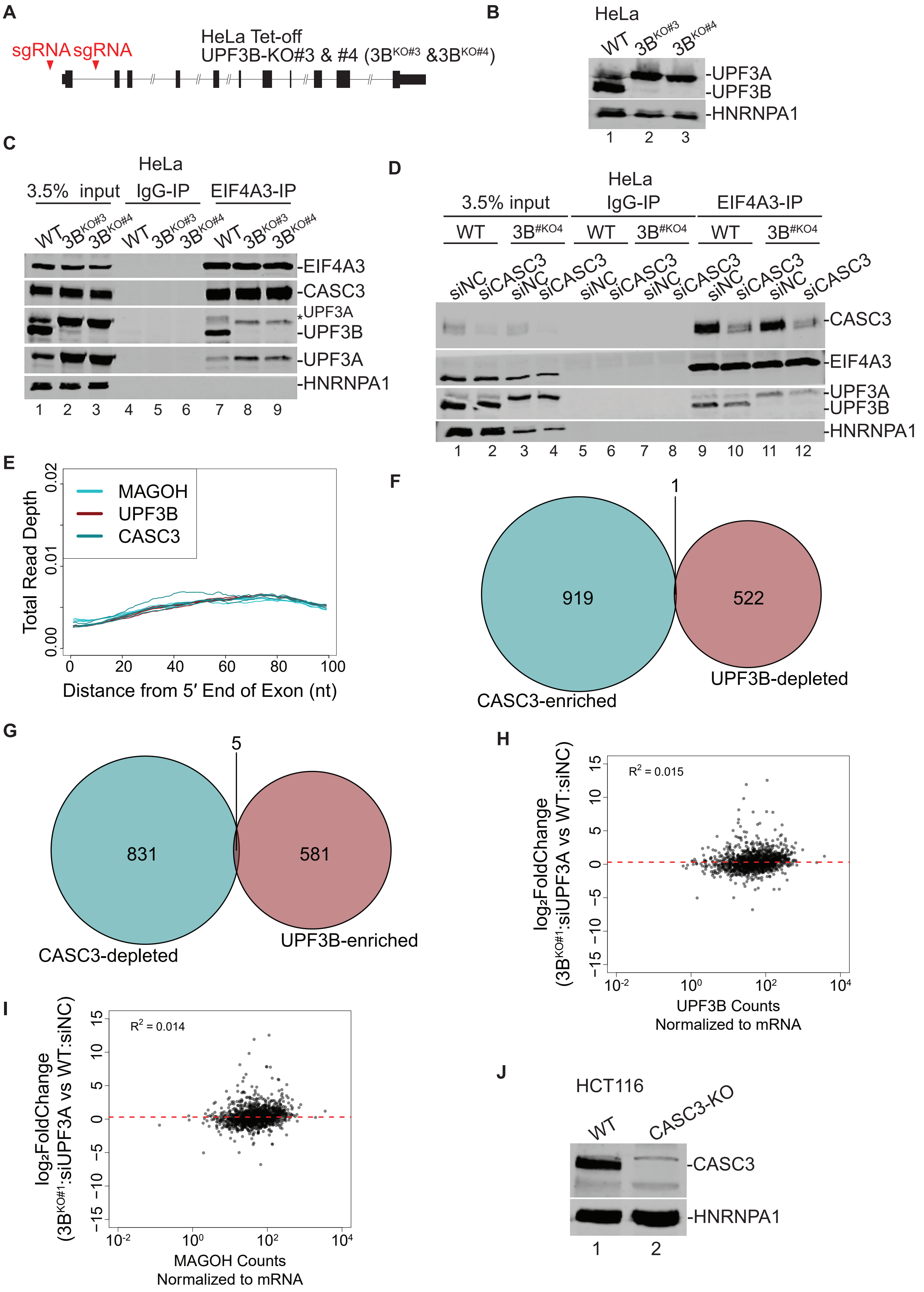


**Figure S3. CASC3 regulates UPF3-dependent NMD.**
A. Schematic of UPF3B knockout in HeLa Tet-off cells using CRISPR-Cas9. Red arrows represent two guide RNA targeting sites which will lead to the deletion of the first exon.
B. Protein immunoblot of UPF3B protein in WT and $3B^{KO}$ HeLa cells. HNRNPA1 is a loading control.
C. Western blots showing proteins on the right in input and EIF4A3-IP from WT and $3B^{KO}$ HeLa cells. Normal rabbit IgG is used for control IP.
D. Western blots as in C from HeLa WT and $3B^{KO}$ cells transfected with either siNC or siCASC3.
E. Meta-exon plot of MAGOH:EIF4A3, UPF3B:EIF4A3, and CASC3:EIF4A3 RIPiT-Seq read-distribution in the 100 nt region from the exon 5′ end.
F, G. Venn diagram of significantly enriched/depleted genes in CASC3:EIF4A3 or UPF3B:EIF4A3 RIPiT-Seq samples as compared to MAGOH:EIF4A3 RIPiT-Seq.
H, I. Scatter plots showing a comparison between gene-level EJC occupancy and NMD efficiency. UPF3B:EIF3A3 (H), or MAGOH:EIF4A3 (I) RIPiT-Seq signal normalized to expression level of individual genes is on the x-axis and NMD efficiency of each gene on the y-axis. For each gene in this analysis, NMD efficiency is the highest fold change (in $3B^{KO\#1}$ siUPF3A to WT siNC) observed for its PTC+ isoform. R2 from the linear regression fit is shown on the top left.
J. Protein immunoblot of CASC3 protein in WT and CASC-KO HeLa cells. HNRNPA1 is  a loading control.

### Figure S4

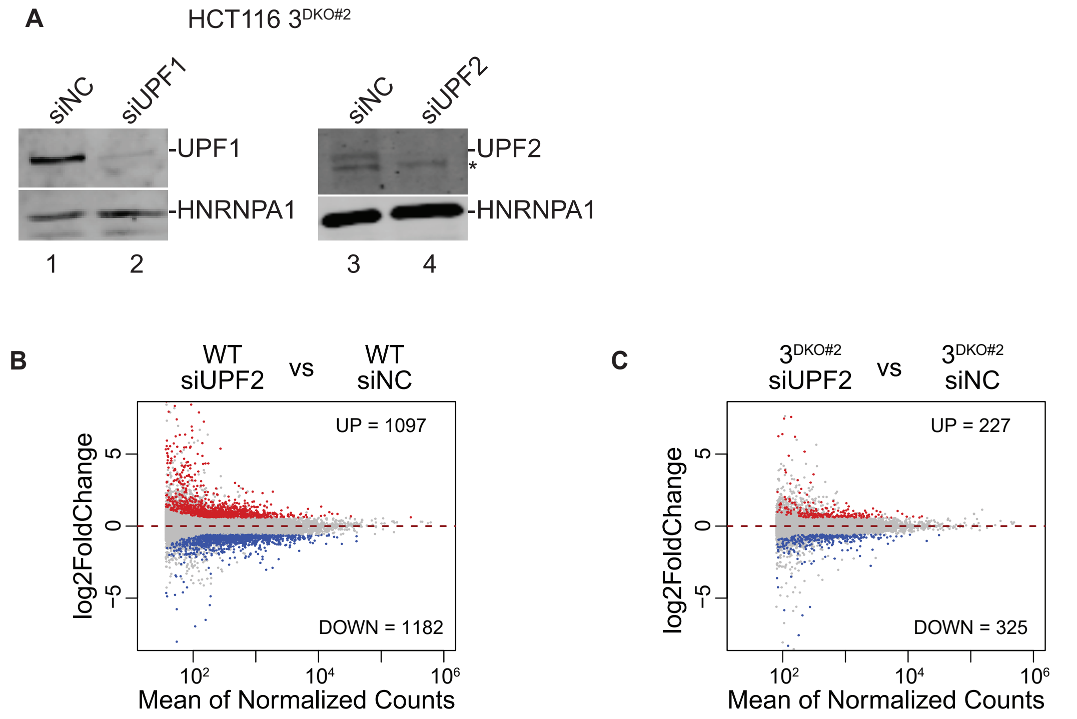


**Figure S4. NMD in the absence of both UPF3 paralogs.**
A. Immunoblots showing levels of UPF1 and UPF2 proteins in $3^{DKO\#2}$ cells that were transfected with negative control (siNC), UPF1-targeting (siUPF1), or UPF2-targeting (siUPF2) siRNAs. HNRNPA1 is a loading control.
B, C. MA plots showing transcript-level changes upon UPF2 (siUPF2) knockdown as compared to control knockdown (siNC) in, (B) WT cells, and (C) $3^{DKO\#2}$ cells. Each dot represents one transcript with average read counts on the x-axis and log2 fold change on the y-axis. Red and blue dots represent transcripts up- or down- regulated more than 1.5-fold with an adjusted p-value < 0.05; these counts are shown on each plot.

### Figure S5

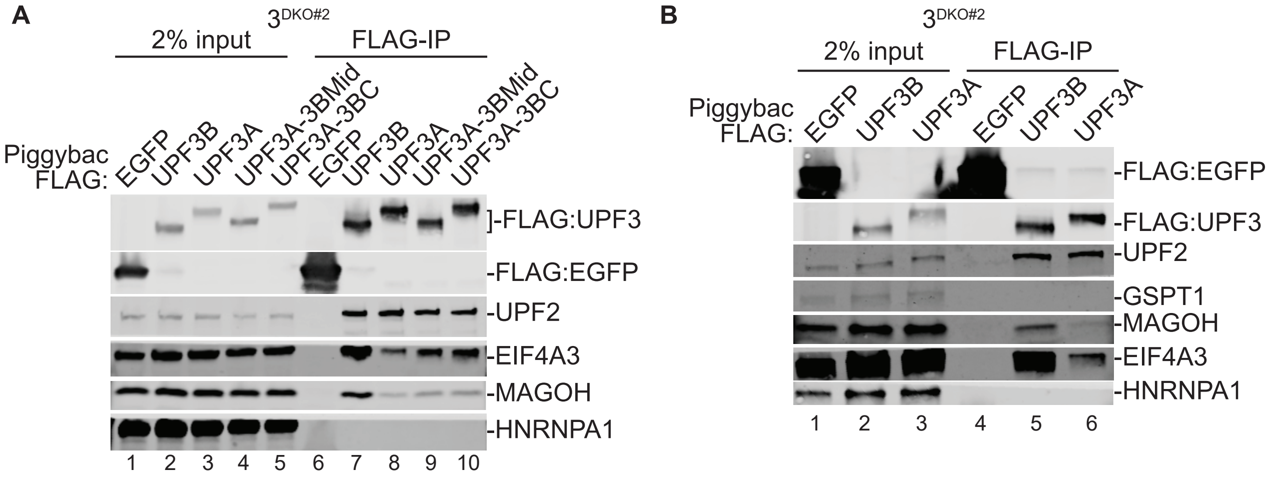


**Figure S5. UPF3 paralogs differ in NMD activity.**
A, B. Protein immunoblots of FLAG-IP from $3^{DKO\#2}$ cells expressing different FLAG-tagged human UPF3 proteins or their chimeras using Tet-on 3G system. HNRNPA1 is used as loading control and RNase A digestion control. FLAG-EGFP is used as an IP control.

### Figure S6

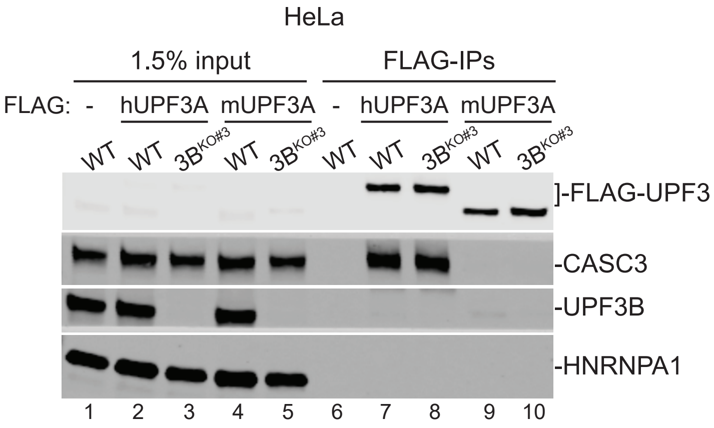


**Figure S6. EJC interaction ability of human and mouse UPF3A proteins.**
Protein immunoblots of input and FLAG-IP from WT and 3$^{DKO\#2}$ cells FLAG-tagged human or mouse UPF3A as indicated above the lanes. HNRNPA1 is a loading and RNase A digestion control.
